# Estimating human mobility in Holocene Western Eurasia with large-scale ancient genomic data

**DOI:** 10.1101/2021.12.20.473345

**Authors:** Clemens Schmid, Stephan Schiffels

## Abstract

The recent increase in openly available ancient human DNA samples allows for new, large-scale meta analysis applications. Trans-generational past human mobility is one of the key aspects that ancient genomics can contribute to, since changes in ancestry – unlike cultural changes seen in the archaeological record – necessarily reflect movements of people. Here we present a new algorithm to quantify past human mobility from large ancient genomic datasets. The key idea of the method is for each individual to compare a hypothetical genetic “origin” point with its actual burial point in space. This is achieved by first creating an interpolated ancestry field through space and time based on Multidimensional scaling and Gaussian process regression, and then using this field to map the ancient individuals into space according to their genetic profile. We apply this new algorithm to a dataset of 3191 aDNA samples with genome-wide data from Western Eurasia in the last 10,000 years and derive a diachronic measure of mobility for subregions in Western, Central, Southern and Eastern Europe. For regions and periods with sufficient data coverage, our mobility estimates show general concordance with previous results, but also reveal new signals of movements beyond the well-known key events.

**Significance Statement:** Ancient human DNA (aDNA) extracted from archaeological contexts (e.g. burials) allows to reconstruct past population movements. Relevant methods work by calculating proportions of shared ancestry among individuals or groups in order to answer specific, regional research questions. Here, we propose a largescale algorithm to quantify human mobility through time and space using bulk aDNA data. The algorithm has two core components: i) Interpolation of the spatio-temporal distribution of genetic ancestry to obtain a continuous ancestry information field and ii) estimation of the spatial origin of each input sample by projecting its ancestry into this field. We apply this to thousands of published genomic samples in the last 10,000 years to trace diachronic mobility patterns across Western Eurasia.

## Introduction

All human behaviour is spatial behaviour and spatial perception and interaction is deeply rooted in the human mind. Individuals have *activity space* [Per+13] and need *personal space* [Høg08]. They interact with each other [ALB18] or the natural environment [Häg05] in space. And, finally, they create entirely new spatial environments [Boi+16]. Understanding movements in space – mobility – on different orders of magnitude is therefore a major component for understanding human behaviour throughout history [Bel14], from the Iceman’s lonely quest through the Ötztal Alps, to the Viking expansion even beyond Medieval Europe, and maybe eventually mankind’s escape to the stars.

Archaeological and anthropological theory provides different concepts and categories to classify mobility. Mobility can be a group property or individual behaviour and it has complex implications for the formation, perception and interaction of identity [Kel92; Ben01; LM15]. Migration, a special type of mobility perhaps to be defined archaeologically as the “*temporal and spatial collimation of a regional source of dispersed cultural property* “[Bur00], is an especially controversial topic [Ant90]. It is notoriously difficult to prove and to uncover its causes among the many interdependencies *micro-* and *macro theories* of migration suggest. Narratives of migration are also notoriously vulnerable to political instrumentalization [Wol19].

The field of archaeogenetics now provides a new perspective on past human mobility and migration, which is at its very core influenced by population genetics theory. The emergence, change and distribution of human ancestry components – mediated by the mobility of their hosts – is in fact one of its most important research questions (e.g. [Haa+15; Lip+18; Fle+19]), causing fruitful and corrective friction with the humanities [Fur18; GF20; Fur21]. While so far much archaeogenetic research focuses on particular cultural-historical contexts, the recent growth of published ancient DNA samples from all around the world enables a new category of quantitative meta analysis.

Large, explicitly spatiotemporal datasets have been part of population genetics research for a long time already [BR19], sometimes even with a focus on mobility quantification [PNS15; BRC16; Al-+19; PPN19]. But to our knowledge up to the time of this writing, only few attempts have been made to systematically derive a continuous, large-scale and diachronic measure of human mobility with ancient genetic data. These are most notably a pioneering publication by Loog et al. 2017 [Loo+17] and another approach by Racimo et al. 2020 [Rac+20]. Loog et al. measure mobility in prehistoric Europe by comparing the distance matrix correlation among spatial, temporal and genetic distance for aDNA samples in moving 4000 year windows. As a result they generate an unscaled mobility proxy curve that indicates elevated levels of mobility correlating to the Neolithic expansion, the Steppe migration and finally the European Iron Age. Racimo et al., on the other hand, employ admixture analysis to individually attribute genomic samples a genetic profile with respect to three specific ancestry components: Mesolithic hunter-gatherers, Neolithic farmers with ancestry originating in the Near East, and Yamnaya steppe herders, arriving in Europe during the third millennium BC. They derive mobility as a wave front speed of surpassed ancestry component thresholds. To overcome sample sparsity and to correlate the arrival of certain ancestry components with biogeographic metrics, they use Gaussian process regression for the interpolation of relative ancestry component occurrence – an idea we also took as a starting point for our proposed mobility estimation method.

In this paper we present a new algorithm (Fig. 1) to estimate past human mobility on the individual level. For each individual it determines multiple spatiotemporal positions of close genetic affinity, which, as we show, can often be interpreted as ancestral origin points. The distance between the location where an individual was buried and these points serves as a proxy for personal mobility in an individual’s (or their immediate ancestors’) lifetimes. We apply this algorithm to several thousand previously published ancient genomes from Western Eurasia dating from between 8000BC and 2000AD (excluding modern genomes) taken from the Allen Ancient DNA Resource (AADR) [Dav21]. And we show that, while the average results largely match expectations including known and large-scale movements at the beginning and end of the Neolithic, these large-scale patterns are accompanied by considerable individual-level heterogeneity.

**Figure 1:**
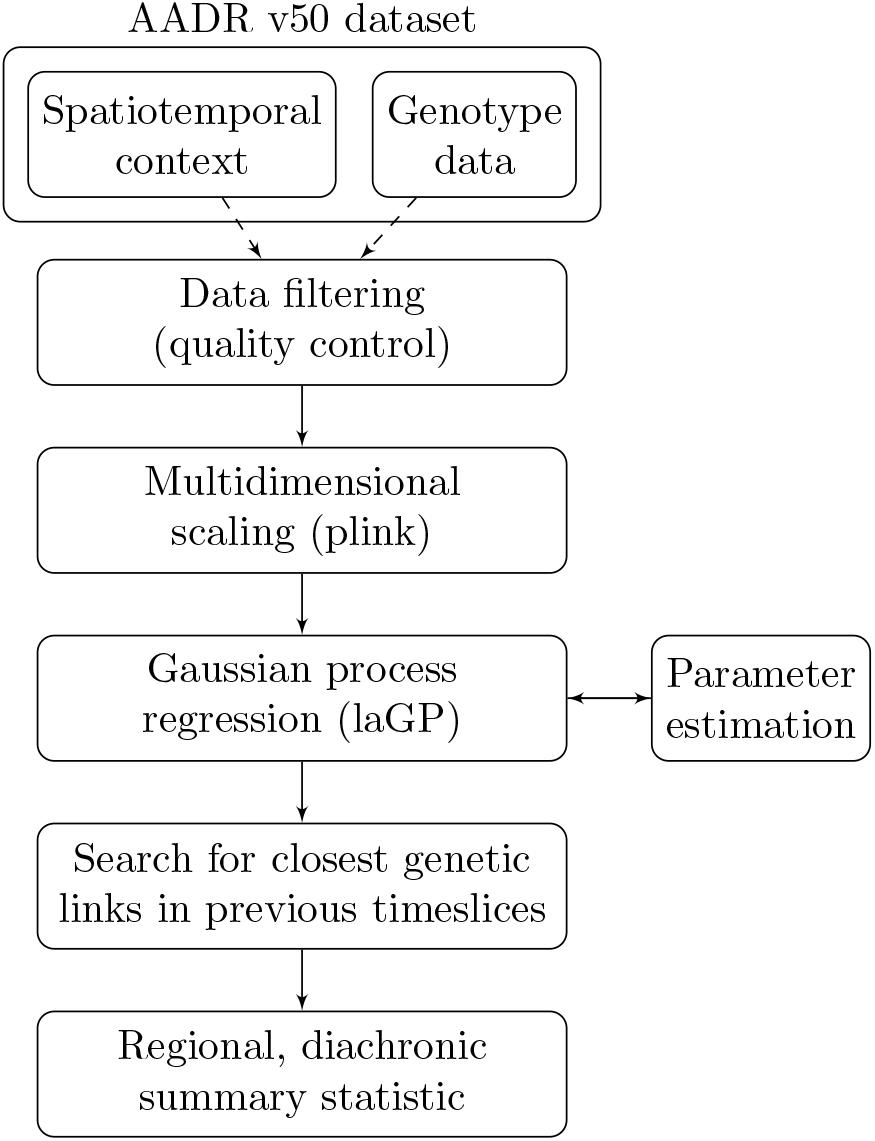
Schematic overview of the general workflow.

## Results

### Interpolating genetic ancestry through space and time

A key challenge for understanding shifts of ancestry through space and time is the inherent sparsity of archaeogenetic data. To counter this we employed an interpolation technique fitted upon 3191 published samples from 85 publications currently available in the AADR [Dav21] for Western Eurasia during the Holocene. All samples in this public data collection reference single-nucleotide polymorphisms (SNPs) from a panel of about 1.24 million known informative positions [Mat+15] and our dataset is filtered according to general sample quality criteria (see Methods). Within the derived subset, the data distribution in time and space is heterogeneous (Fig. 2), with generally few data points from the European Mesolithic, significantly more from the Neolithic, then most from the Bronze Age and again less from the Iron Age and later periods. The diachronic amount of data from Great Britain, Iberia, Central Europe and Southeastern Europe is comparatively high, whereas other regions are less well covered.

**Figure 2:**
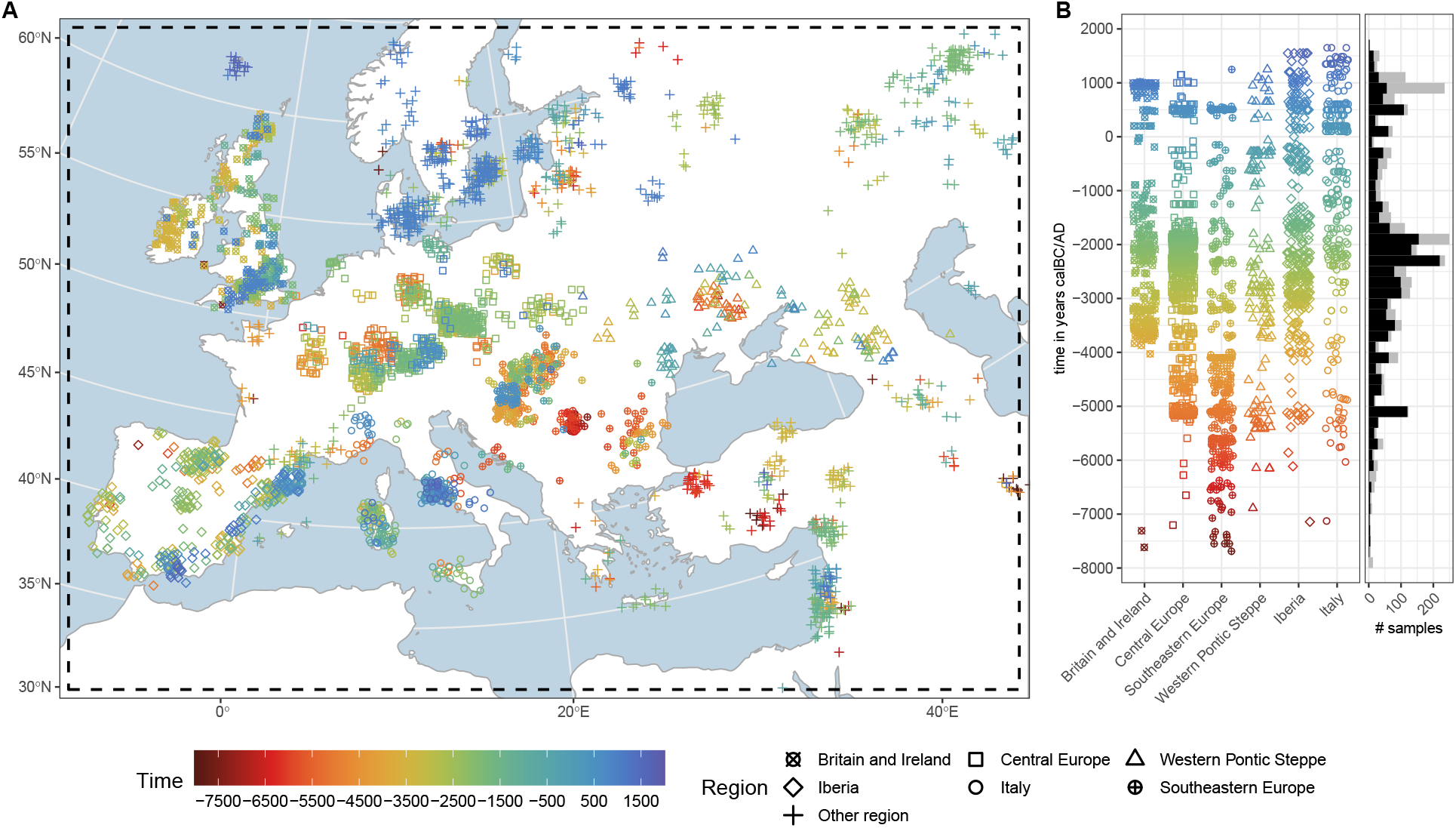
Spatial and temporal distribution of the aDNA sample selection. **A**: Map (EPSG:3035, ETRS89 Lambert Azimuthal Equal-Area, “European grid”) with the research area (dashed). Samples are jittered with up to ±60km in x and y direction to reduce the effect of overplotting. The sample dots are coloured according to their temporal origin, where the time is given in years calBC/AD (negative values indicate ages calBC). The sample dot shape encodes the attribution to different analysis regions. **A**: Scatter plot of temporal sample distribution for each analysis region. The stacked histogram on the right shows the sample count through time for all samples in grey, and for the ones within the defined analysis regions in black (bin width = 200 years).

We applied multidimensional scaling (MDS) on this data to reduce its dimensionality to two summarising ancestry components (Fig. 3), that are by construction most informative about the genetic diversity across the sample set (see Supplementary Text 3.1 and Supp. Fig. 7 for a run with three result dimensions). We find the largest internal separation of samples to be along the tempo-cultural boundary between the Mesolithic and the Neolithic, highlighting the strong population shift the Neolithic introduced into Europe [Laz+14; Haa+10; Sko+12]. Many other patterns seen in the MDS are also consistent with previous observations and will be discussed among our results below.

**Figure 3:**
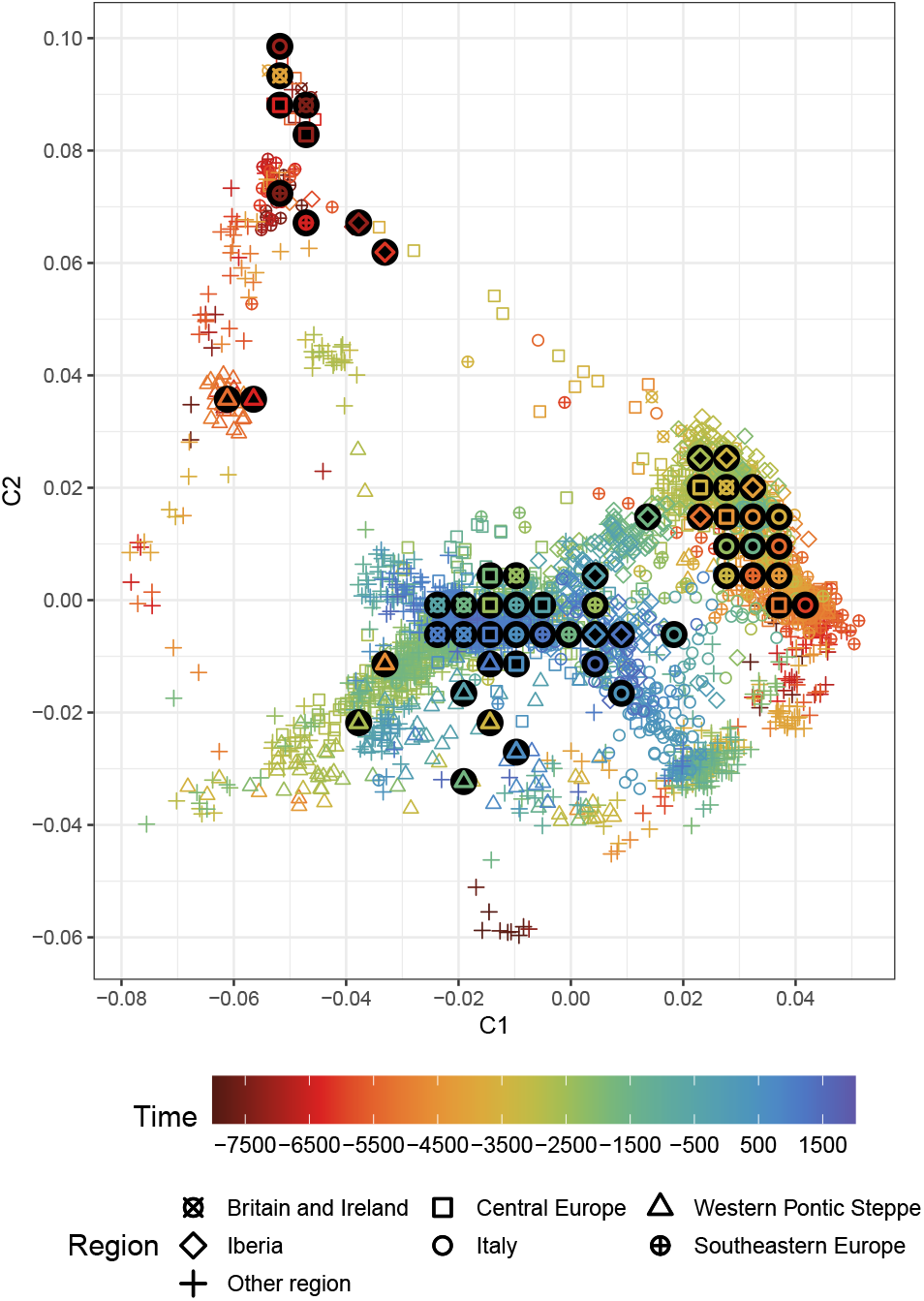
Scatter plot of the sample distribution in 2D multidimensional scaling space. Each sample is plotted with the same shape and colour as in Fig. 2. The bigger, black circles are the centroids of region-time groups (bin width = 1000 years). To prevent overplotting, the centroids are not printed on their exact positions, but instead rearranged in a non-overlapping lattice. See Supp. Fig. 7 for a 3D version and Supp. Fig. 8 for a larger version where the individuals mentioned in the text are highlighted.

We modelled these two MDS-derived ancestry components individually as the dependent variable in a Gaussian process regression (GPR) model with three independent input variables describing the position of each sample in space and time. To learn the properties of the relevant covariance matrix (*“kernel”*) for a model with the best mean postdiction abilities we explored multiple methods: Variogram analysis, maximum likelihood estimation and cross-validation (see Supplementary Text 1). With the parameterized Gaussian process regression model we predicted iterations of average spatio-temporal genetic ancestry grids across Europe (e.g. Fig. 4), explicitly sampling from the the temporal uncertainty of the input data.

**Figure 4:**
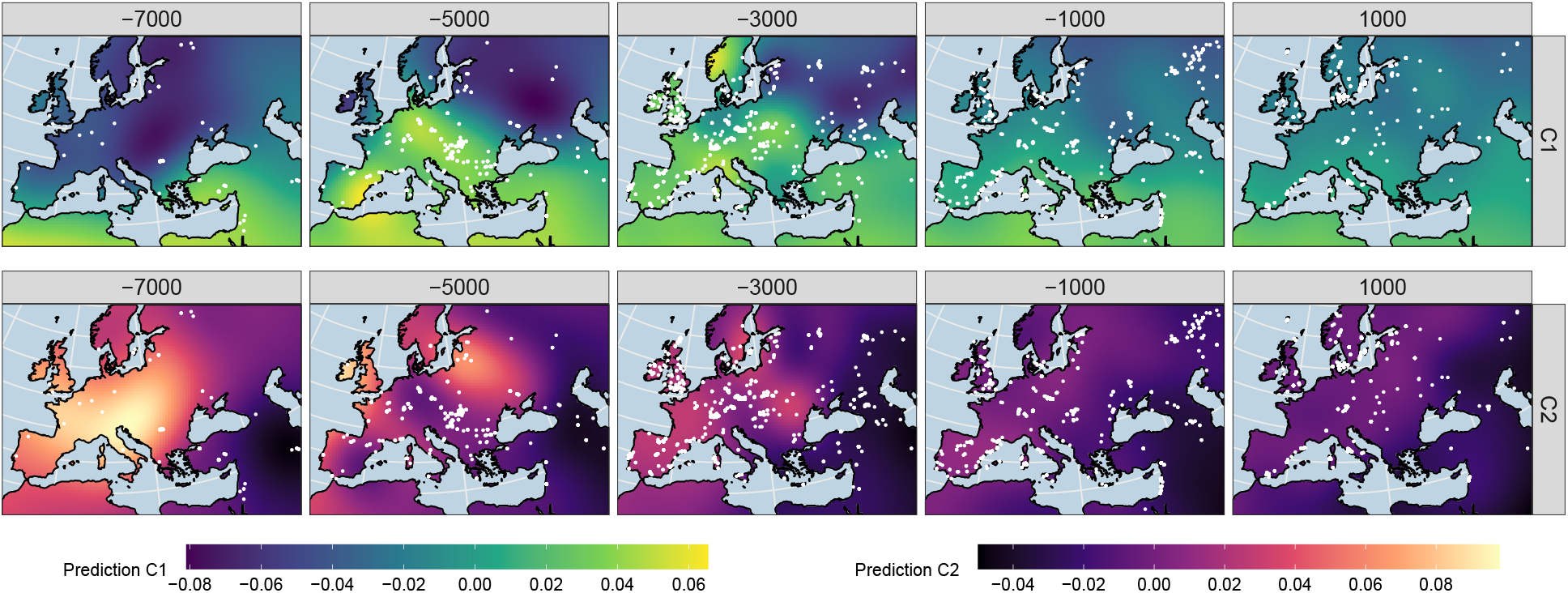
Gaussian process regression interpolation map matrix based on the multidimensional scaling dimensions (resolution: 50km). The five maps on top show timeslices through the interpolated, spatio-temporal 3D space for the derived ancestry component C1, the five on the bottom for C2. The samples from a time window *±*1000 years around the slicing position are plotted as white dots.

The interpolation of ancestry components across time and space reflects how 10 millennia of human population changes have shaped genetic ancestry in this area. As already seen in Fig. 3, both ancestry components C1 and C2 here most strongly reflect the enormous changes that underlay the transitions during the Early Neolithic, with increasing values (for C1 coloured in yellow in Fig. 4) throughout Central and Western Europe around 5000BC as a result of people moving north-westwards from the Levant and western Anatolia. They also prominently feature further changes after 3000BC, bringing ancestry previously located in Eastern Europe and the Eurasian steppes into Western and Central Europe.

With this interpolation one can attempt the reconstruction of continuous, local ancestry histories even for places without consistent data coverage. To illustrate this, we selected arbitrary spatial positions (corresponding to six capital cities) and used the GPR model to postdict how the genetic profile in these locations changed through time (Supp. Fig. 4). The six “virtual” time-series again generally reflect our knowledge of the genetic changes in Europe: In London, Budapest, Rome and Barcelona we observe an ancestry shift with the arrival of Neolithic, and then once more with Steppe ancestry – with small regional differences. Riga, on the other hand, starts out with a higher degree of Eastern Hunter-Gatherer (EHG) ancestry before skipping the influx of the Anatolian farmer component. Jerusalem, expectedly, fills a markedly different spot on the genetic map.

### Estimating individual-based mobility

While the interpolated ancestry field reflects the *average* change in ancestry through space and time, it also forms the basis for our proposed algorithm to understand *individual* -based mobility. The key idea is to assign to each individual a hypothesised place of closest genetic affiliation in a previous field timeslice (i.e. multiple hundred years before said individual), which we interpret as point of ancestral origin. The difference of this ancestral origin point and the factual burial position is a quantitative measure of ancestry relocation through inter- or intragenerational mobility. As an example, consider an individual from Roman-time Britain (individual 3DRIF-26 from ref. [Mar+16]), buried in York, but featuring a genetic ancestry profile from the Near East. This is a clear case where the authors concluded that either this individual themselves or their ancestors came from the Near East, but ended up in Britain. In this case, we would thus infer an “origin- or mobility vector” pointing from Britain to the hypothesised region in the Levant, resulting in an origin-distance of several thousand kilometres and in south-western direction (Fig. 5C).

**Figure 5:**
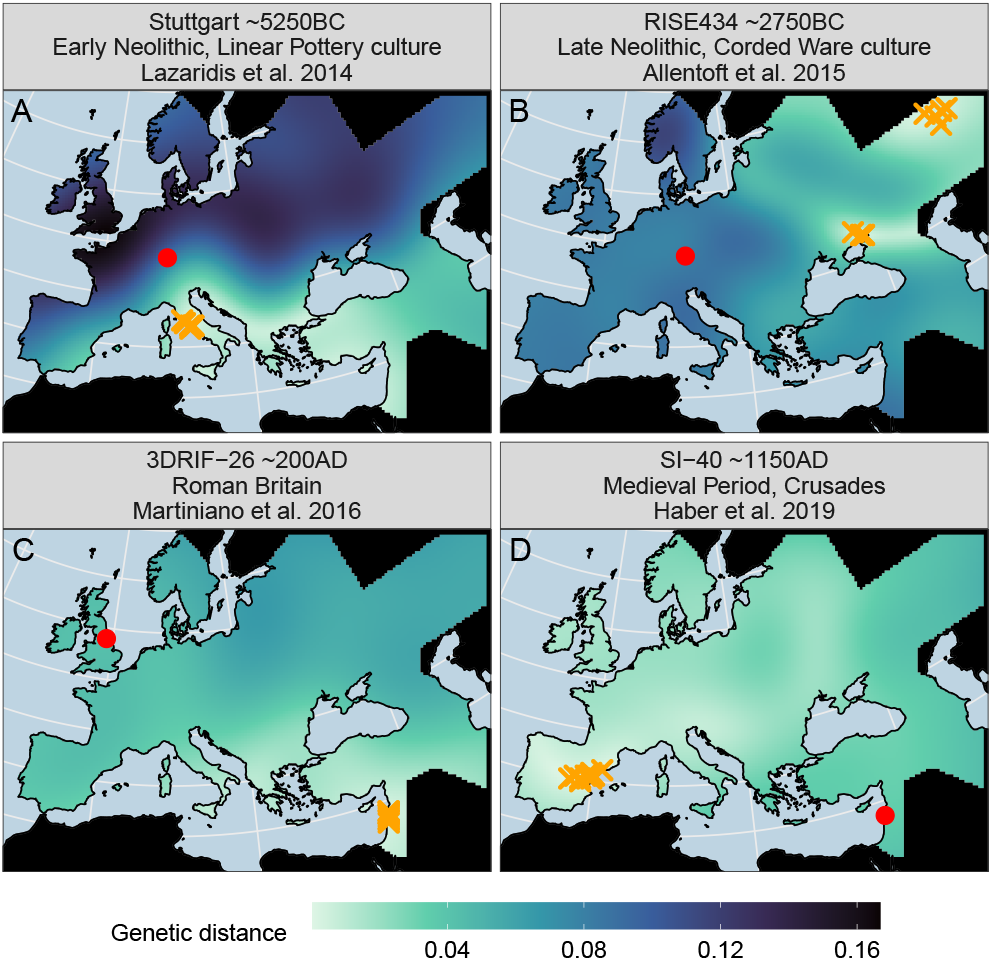
Genetic distance map matrix for four selected individuals. The red dots show their burial position. For each individual ten temporal re-sampling iterations of prediction grids (resolution: 30km) for C1 and C2 were created, where each grid is a timeslices several hundred years before the median burial age – just as for the slices in Figure 4. For these timeslices the genetic distances (Euclidean distance in 2D MDS space) between the individual and each field grid point were calculated, thus creating a smooth raster indicating regions with increased or decreased genetic similarity to the respective individual. The orange crosses mark the ten points of minimal genetic distance.

It is indeed instructive to consider our approach with other well understood individuals that represent outliers in their genetic signatures, and which have been used in the past to illustrate mobility. Figure 5 shows four such concrete cases for specific published samples. The individual named Stuttgart, one of the first ancient genomes sequenced [Laz+14], is also one of the earliest Neolithic samples from Central Europe. They display non-local genetic ancestry in the sense that they differ strongly from preceding Mesolithic samples in the area. In our analysis, we show that indeed the lowest genetic distances for this individual can be found in Anatolia and Southeastern Europe (Fig. 5A). This indicates mobility from there to Central Europe in accordance with archaeological models [Por+20]. The lowest genetic distances can be observed to Southern Europe, where the Neolithic expansion followed another route [Boc+09; Boc+12]. In the late Neolithic, individuals affiliated with the Corded Ware culture from Central Europe have been identified as one of the earliest with so-called Steppe ancestry, which was present already before 3000BC in the Pontic Caspian steppe. Indeed, for a representative sample from that group (RISE434), we find the closest matching ancestry points falling into Eastern Europe (Fig. 5B). Finally, confirming the analysis by Haber et al. 2019 [Hab+19], we find that multiple samples (for example SI-40) extracted from a mass burial near a Crusader castle in Sidon in present-day Lebanon are linked to Iberian ancestry profiles (before the Umayyad conquest) (Fig. 5D).

### Regional mobility patterns during the last 10,000 years in Western Eurasia

Given our interpolation model for the average ancestry through space and time, and the individual-based mobility as computed using the origin search algorithm described above, we can derive a mobility-vector for every single sample in our dataset and investigate these vectors through time. The results of that analysis are summarised in Figure 6, both in terms of the lengths of individual mobility vectors (shown on the y-axis) and direction (shown in colour according to the legend). While in principle we can apply our algorithm to every sample in the dataset, we here focus on a selection of confined regions with acceptable coverage of samples throughout the study time period (Fig. 2). We mostly consider patterns emerging from long-distance signals, observed as individual points with large origin-distances (typically 1000km and further), as these typically correspond to events described previously in the literature and thereby provide a proof-of-concept for our method. However, beyond these long-distance signals, we highlight a considerable level of complexity of smaller-scale signals that may correspond to previously unknown events. Shown along the individual distances is a moving average curve together with an error band (in grey shading), which in some cases may help putting the largest individual-based events into context. Alternative visualizations of the time series shown in Figure 6 are available with the Supplementary Figures 5 and 6.

**Figure 6:**
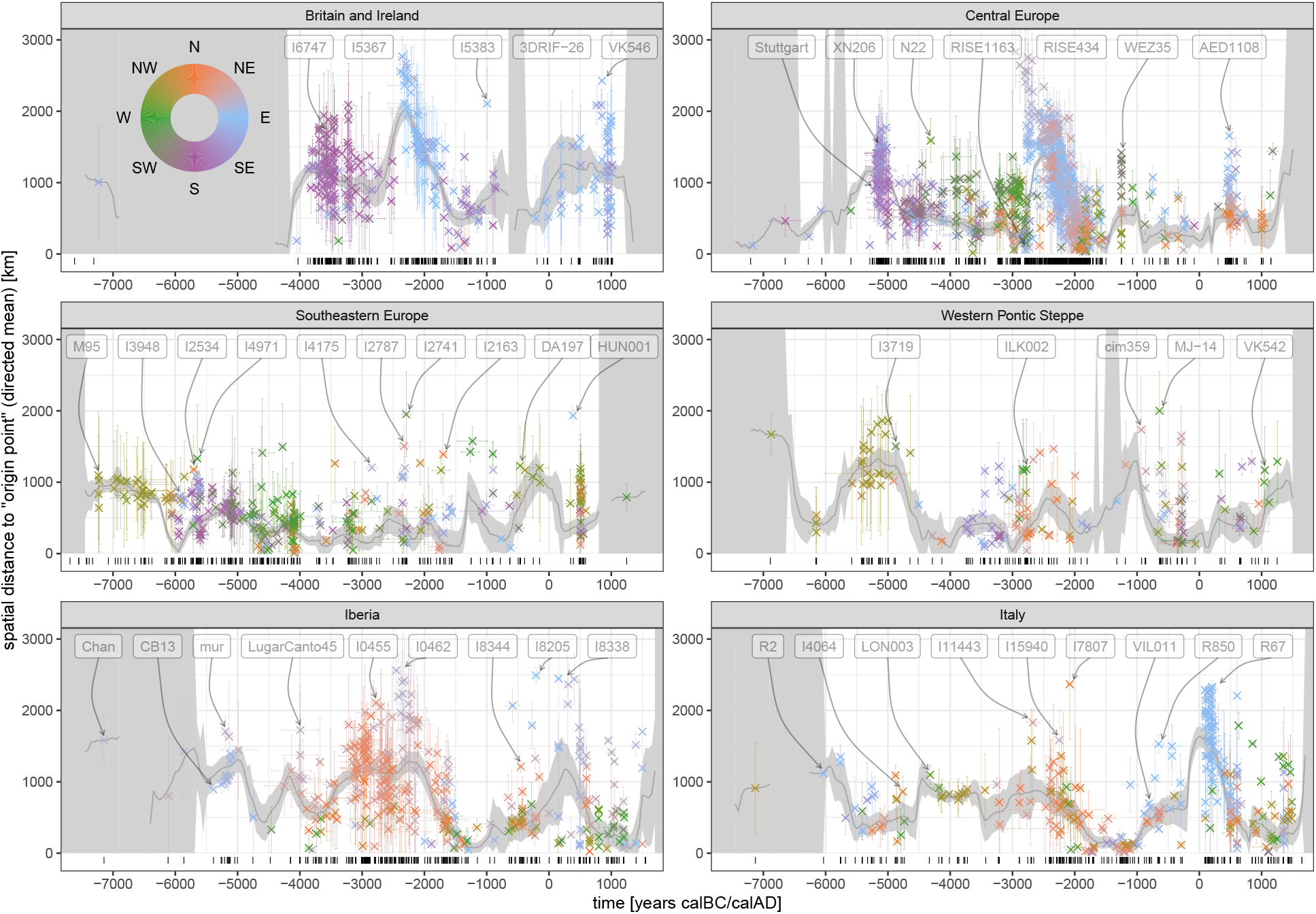
Mobility estimation results. One scatter plot for each of the six analysis regions (see Fig. 2). Each dot represents the mean mobility vector for one sample across 100 temporal re-sampling runs. The position on the x-axis is the mean of the sampled ages in years calBC/AD, and the position on the y-axis is the mean distance to the 100 origin points, with dot colour encoding the mean direction according to the circular legend in the top-left corner. Each observation comes with error bars on the x- and y-axis, covering one standard deviation of the 100 resampling observations. The distance mean per observation is a directional mean, so opposing vectors cancel each other out, whereas the standard error is calculated from absolute distances. The barcode plot at the bottom of each subplot documents the diachronic data coverage for each region. The smooth grey curve printed below the samples is a 400 year moving mean for the spatial distance. It is calculated from the total set of the 100 resampling iterations. The dark grey ribbon accommodating this mean curve, on the other hand, is the standard error of the mean based on the mean dots displayed here. It is visualized as infinite if a given 400 year time window has less than three observation.

Beginning with our time series from Great Britain and Ireland, the largest observed individual signals correspond to the Early Neolithic in the 4th millennium BC [She10; Tho15; Bra+19; Sán+19a; Cas+20], the Bell-Beaker transition after around 2500BC [Cas+16; Ola+18], Roman Britain and the Viking period [Mar+20b]. Note for example the indicative individuals I6747 [Bra+19], I5367 [Ola+18], 3DRIF-26 (*>*3000km; already discussed above) [Mar+16] and VK546 [Mar+20b], which each represent extremely long distance mobility. Direction-wise, the respective mobility peaks are consistent with what we know about the sources for these events, with Southern sources during the Neolithic, and Eastern sources during the Bell-Beaker and Viking periods. The Neolithic transition is visible in our mobility proxy as a clear upwards-jump, both in the average and individual origin distances, contrasting the few sufficiently well covered and apparently very “local” Mesolithic individuals from before 4000BC. But this peak only dies down surprisingly slowly until the first half of the third millennium, attributing almost every Neolithic individual a foreign “origin”. To some degree this might be an effect of the smooth, only slowly recoiling ancestry interpolation model and the peripheral position of Britain and Ireland, which renders origin positions on the continent disproportionately likely. But the tardiness of the recovery also supports the assumption of a large and stable sphere of interaction or at least strong genetic similarity across Western Europe during the Neolithic. We will discuss a corresponding observation below for Central Europe and Iberia. The following Bronze Age peak, triggered by incoming ancestry ultimately from Eastern Europe, is remarkably strong (see also Supp. Fig. 6) and persists even after the initial Bell-Beaker transition. For the period after 2000BC Olalde et al. 2018 suggest a much more homogeneous gene pool, which does not rule out the possibility of “incoming continental populations with higher proportions of Neolithic-related ancestry” [Ola+18], though. These might be one reason for the mobility vectors pointing to the East and South in the Middle Bronze Age. In the Late Bronze Age the individual I5383 [Ola+18] stands out with a long mobility vector pointing to Eastern Europe. As they are not separated from their contemporary peers in MDS space (Supp. Fig. 8), we assume their outlier position to appear incorrectly inflated here.

For Central Europe we observe similar peaks as for Britain and Ireland. The Neolithic expansion reaches this area in the late sixth millennium and leads to a first, strong uptick of the mobility signal from the Southeast [GD15; Lip+17; Nik+19; Bru+20], visible for example in the aforementioned Stuttgart individual [Laz+14] or XN206 from Stuttgart-Mühlhausen [Riv+20]. These individuals’ origin points cover a wide corridor from Western Anatolia to the Balkans, with some directed also towards the Southern route of the Neolithic in Italy. Not unlike the post-Neolithisation development in Britain, the mobility pulse then dies down only slowly in the fifth millennium. An interesting case in this spatiotemporal context is individual N22 from modern day Poland. Fernandes et al. 2018 [Fer+18] describe them as “the most recent individual (*≈*4300 BCE) with a complete genomic WHG attribution to be found to date in an area occupied by Danubian Neolithic farmers”, which causes our origin search to link them to remaining hunter-gatherer populations i. a. in Ireland. In the first half of the third millennium Steppe ancestry arrives, as observed in many Late Neolithic Corded Ware individuals – like aforementioned RISE434 [All+15]. Their origin vectors then clearly point into the far East and Northeast [Haa+15; All+15; Bru+20; Fur+20; Lin+20]. This strong signal, lasting well into the Bronze Age, is heterogeneous both in distance and directionality. We caution that the spread of Steppe ancestry did most likely not follow a perfect wave-of-advance-like pattern, leaving pockets of unaffected or only later-affected ancestry behind, which will inevitably result in more erratic mobility estimates. After 1500BC the data density for Central Europe decreases and general observations become more difficult. Given archaeological and eventually historical evidence, it is of course not unreasonable to assume a high degree of mobility in the Late Bronze Age, the Iron Age [KL05], and the Medieval period, connecting Central Europe to France, Great Britain, Southern Scandinavia, Eastern Europe and the Balkans, catalysed by different cultural processes [Ola+18; Mit+19]. Two remarkable individuals with long mobility vectors are WEZ35, which is representative for the relatively unstructured population documented from the Tollense Bronze Age battlefield in northern Germany [Bur+20], and AED1108 from Bavaria with strong skull deformation and about 20% East Asian ancestry [Vee+18].

Southeastern Europe stands out in our analysis, because we could include a high number of comparatively early samples from the Mesolithic, e.g. M95 [Gon+17]. All of them are from the small and extraordinary Iron Gates area in Serbia and Romania, where the Danube passes between the Balkan Mountains and the Southern Carpathian Mountains range. They therefore may not be representative for the entire region and their long mobility vectors pointing to the Northwest are probably an artefact of the also low data density in Central Europe and Britain, where our ancestry field misrepresents a most likely more Western Hunter-Gatherer-related (WHG) profile. During the 6th millennium BC, correlating well with the beginning of the Neolithisation in the region [Por+20], we observe more plausible non-locality signals: The ancestry profile of Early Neolithic individuals like e.g. I3948 [Mat+18] from the Adriatic coast, points, to Western Anatolia. Surprisingly, for the individuals I2534 and I4971 [Mat+18; Lip+17], we observe a large origin distance to the North and West, even after the onset of the Neolithic. These individuals might not have been (personally or through his immediate ancestors) part of any permanent long-range mobility: they lived at a time and place where new ancestry was arriving with the Neolithic package – rapidly changing the local ancestry landscape – and their genetic “displacement” thus becomes an indirect proxy of the major mobility event surrounding them. I4971 even features some “first European farmers” (FEF) ancestry [Lip+17], which positions them between a local hunter-gatherer and the new farmer profile (Supp. Fig. 8). Unlike other European regions, the arrival of Steppe ancestry in Southeastern Europe appears more gradual, beginning earlier and less abruptly [Mat+18]. Few individuals show a clear mobility signal pointing to the far Northeast – e.g. I4175 [Mat+18]. For later periods, finally, we observe some remarkable outliers with strong mobility signals: For example the Hungarian Bell Beaker individuals I2787 and I2741 [Ola+18], the Middle Bronze Age individual I2163 from Bulgaria [Mat+18], the Iron Age Scythian DA197 [Bar+18] or the Migration Period Hunnic individual HUN001 [Gne+21].

Even further to the East, in the Western Pontic steppe (including the area north of the Greater Caucasus mountain range), we see a quite varied account of ancestry influx. For the Ukrainian individuals from the sixth millennium and before, Mathieson et al. 2018 [Mat+18] report ancestry on a cline between Eastern-, Scandinavian Mesolithic- and later Western Hunter-Gatherers. This genetic affinity is reflected in the first increase of signal we observe mainly from the Northwest during the sixth millennium, confirming previously described similarities in the developments in Eastern and Northeastern Europe [Jon+17]. Only at the beginning of the fifth millennium one extraordinary Individual (I3719) stands out with “entirely northwestern-Anatolian-Neolithic-related ancestry” [Mat+18] and thus long-distance affinity to the West and Southwest. Most data for the Neolithic and the Bronze Age is from the Caucasus region and documents a complex, though relatively local mobility history [Wan+19]. Within this time frame, multiple Globular Amphora context individuals (e.g. ILK002 [Mat+18]) from present-day Ukraine stand out with a strong origin signal from the West. During the Iron Age, more individuals with a relatively long-distance mobility signal appear, most notably cim359 [Krz+18] and MJ-14 [Jär+19]. Their origin vectors point to the opposite ends of Europe, illustrating the region’s position as a bridge between Europe and Central Eurasia housing different equestrian steppe nomad populations – e.g. Cimmerians, Scythians, and Sarmatians (see also Supp. Fig. 5). This generally holds true into historical times, including the Migration-[Bar+18] and Medieval Periods (e.g. VK542 [Mar+20b]).

Already the first hunter-gatherer individual available from Iberia – Chan [Gon+17] – has a very large mean mobility vector. This signal is not reliable, though, given the fact that no local, preceding reference data exists, which could inform the ancestry field for the origin search to appreciable accuracy. Much more relevant are the observations for Early Neolithic individuals like CB13 [Ola+15] or mur [Val+18]. They document the southern route of the Neolithic expansion. From the second half of the fifth millennium to the middle of the third millennium many individuals from Iberia are attributed long origin vectors towards the North and Northeast (e.g. LugarCanto45 [Mar+17] or I0455 [Ola+18]), although others have described this period as a time of relative genetic stability [Ola+19]. This forms a parallel observation to Southern- and Western-facing origin vectors described above for Great Britain, Ireland and Central Europe between 4000 and 2500BC. We suspect that this crisscrossing of origin vectors may be caused by the low levels of genetic differentiation among different Neolithic populations. The Neolithic expansion and the following resurgence of hunter-gatherer ancestry in populations in Iberia, Central Europe and Great Britain might have created a large geographic area of very similar genetic ancestry (see also Fig. 3). Alternatively – or additionally – the Atlantic sphere of influence connecting Western European megalithic cultures (to be taken up later in the Bell Beaker phenomenon and beyond) could have indeed induced a high degree of mobility in said region [Tho15; Pau19; Sán+19b]. More clearly interpretable signals emerge later in the third millennium in Iberia with the arrival of Steppe ancestry – well visible through origin vectors pointing to the far Northeast for individuals like I0462 [Ola+18]. In the Iron Age and later we observe some non-locality from the North (e.g. I8344 [Ola+19]) – which could potentially be connected to the spread of Celtic languages to the region [Fis+18] – but then mostly from the East (e.g. I8205 and I8338 [Ola+19]), possibly through Greek and Roman influence. We note that the relative lack of samples from Northern Africa forced us to mostly exclude it from the genetic similarity search in our analysis, which masks all potential mobility that might have taken place between Europe and Africa [Gon+19].

The final focal region studied here, Italy, comprises not only the Italian Peninsula, but also Sicily and Sardinia. These go through partially independent developments not comprehensively represented in the available data. Samples from the sixth millennium are limited to Sicily as well as Northern and Central Italy. They fit well to what we know about the southern route of the Neolithic Expansion with ancestry arriving from the East [Ant+19]. Indeed, the ancestry vectors of Early Neolithic samples like R2 [Ant+19] point directly to Western Anatolia. A few hundred years later the Neolithic ancestry profile is distributed across large parts of Europe and our derived mobility proxy reflects less a point of origin for the respective Neolithic samples, but rather their entanglement in the preceding cross-European mobility phenomenon. We assume this to be the reason for the moderately strong mobility signal we measure from the the fifth to the middle of the fourth millennium (e.g. I4064 [Fer+20] and LON003 [Mar+20a]), arising despite almost all our input data is from Sardinia, where others have observed genetic continuity until the first millennium BC [Mar+20a]. In the third millennium Steppe ancestry arrived on the Italian peninsula, heralding multiple long-distance mobility signals: The affected Sicilian and mainland individuals show affinity to the North and East – most notably I11443 [Fer+20] from Sicily, which was reported to have the highest amount of Steppe ancestry in ref. [Fer+20]. An even more extreme outlier, I7807, who has no Steppe ancestry, is misplaced by our algorithm, likely because absence of Steppe ancestry is rare at this late time in Europe. Their genetic profile thus lacks sufficient spatial support in our model and our search gets pushed towards the fringe of our analysis region. Note the Chalcolithic sample I15940 [Fer+20] from Sardinia with their eastern mobility signal. Fernandes et al. 2020 identified them as an outlier with “significant affinity to Levantine and North African Neolithic individuals”. The second millennium in our Italy timeseries is almost exclusively covered by samples from Sardinia and Sicily, with a low mobility proxy signalling genetic isolation. During the Iron Age, Sardinia and the Italian mainland become once more part of an exuberant Mediterranean mobility network, as shown for example by the relatively long mobility vectors for VIL011 from a Carthaginian/Phoenician-Punic context [Mar+20a] or R850 [Ant+19], which Antonio et al. 2019 could model as a “mixture between local people and an ancient Near Eastern population (best approximated by Bronze Age Armenian or Iron Age Anatolian […])”. Given North-Africa was excluded from the origin search, many signal from this time period are not entirely reliable and miss an important ancestry component. We finally observe the most extreme signals of non-locality in Italy during the height of the Roman empire, in the first centuries AD, where a unique pattern of mostly Eastern non-local origin emerges, consistent with strong Near Eastern influx into the city of Rome (visible e.g. with individuals like R67 [Ant+19]).

## Discussion

The method to estimate human mobility from genetic data we presented here is based on a simple key principle: Changes in genetic profiles are informative about population movements. This key principle is not new, and in fact is the core assumption behind all major archaeogenetic studies that have revealed mobility in the past. Most notably in Western Eurasia movements associated with the Neolithic expansion (e.g. [Haa+10; Sko+12; Ola+15; Lip+17; Fer+18; Fre+18; Bra+19; Nik+19; Riv+20]) and the arrival of Steppe ancestry (e.g. [All+15; Haa+15; Cas+16; Ola+18; Mit+19; Fer+20; Fur+20; Lin+20]). In our algorithm, we have used this basic principle to derive mobility at an individual level, by interpreting genetic profiles as quantitative proxies for a biogeographic field. A key challenge is the fact that the nature of this genetic-spatial mapping changes through time, due to human movement, the very subject of this study. Conceptually, there can never be a perfect solution to this challenge, since ultimately genetic ancestry is *not* tied to geographic space, but to mobile people living in this space. In our method, we have addressed this by using Gaussian process regression to approximate an *average* ancestry field, which is by construction forced to change only slowly through space and time. It thereby forms the basis to determine individual-wise mobility as differences in relation to the slower group-based shifts in ancestry. These choices result in individual and average mobility signals which generally fit the published state of research as reconstructed from the very same samples. The Neolithic demographic expansion, the Steppe migration and a number of smaller population turnover events show clearly visible signals for most of our study regions.

An interesting result of our analysis concerns the role of outliers: In many described events of “massive” scale, the underlying signals are most visible in the presence of multiple individuals with very long origin-distances according to our algorithm. Indeed, we expect our mobility estimation to perform well in picking up outlier individuals who moved over a long distance in a short amount of time. The smaller the spatial distance of a mobility event and the longer the duration of the process, the more diffuse and unclear the respective signal gets, always highlighting pioneers over latecomers. But at the same time, we observe that there are many individuals with arguably more “local” genetic signatures. A possible prospect for future development would be to estimate fractions of migrants among groups, to gain a more nuanced understanding of how mobility affected communities. Significant shifts in local ancestry proportions can not only be the outcome of the often cited deliberate “mass migration”, but potentially also of bottlenecks [Lin+16], forced migration [Mic+20], changes in reproductive behaviour [SP07], or the influence of a highly fertile minority [Bal+15].

To improve the results obtained in this paper, several important directions may be taken. Future research will probably be in a position to include more data as the sampling gaps in world wide ancient DNA data are quickly filled. That will make large-scale meta analysis more and more feasible and will allow for increased postdiction model resolution. Beyond that, developing more sophisticated spatiotemporal interpolation models will be a core challenge. We are convinced that Gaussian process regression is a very powerful method, but maybe other approaches allow for more heterogeneous covariance settings dependent on the data density in space and time, or even involve full-scale machine-learning [BRK20]. One challenging aspect of an interpolation through space and time, like in our method, is that it is not able to capture situations of multiple co-existing ancestries living in close proximity. Interpolation will in such cases create an average ancestry profile which may not be meaningful. Also for the mobility estimation method entirely different algorithms may be conceived, to get a more robust and precise measure compared to the one we present here. It may, for example, be possible to assign priors for the search of ancestry “origin” points and not follow genetic information blindly. This could include mobility information from artefact refitting [Clo00], isotope analysis [Bri20], least-cost-path calculation [VNG19] or genetic kinship analysis (e.g. [Mit+19]) – as derived by archaeology or other neighbouring disciplines. Similarly, it may be possible to codify linguistic, historical or even archaeological data to derive alternative, quantitative large-scale measures of human mobility [RHS19].

## Materials and Methods

All code for this paper with all relevant scripts and data is available in a repository here: https://github.com/nevrome/mobest.analysis.2022 (an archived version with DOI will be added upon publication). From that we outsourced the main mobility estimation workflow into an R package available here: https://github.com/nevrome/mobest. All data analysis and plotting was done in R [R C21] with the following packages: Bchron [HP08], checkmate [Lan17], cowplot [Wil19], data.table [DS19], DiagrammeR [Ian20], fields [Dou+17], fractional [Ven16], future [Ben21], ggrepel [Slo21], ggridges [Wil21], igraph [CN06], khroma [Fre21], laGP [Gra16], latex2exp [Mes15], lemon [Edw20], raster [Hij21], rnaturalearth [Sou21], sf [Peb18], viridis [Gar+21] and finally the tidyverse and the many packages within it [Wic+19].

### Dataset

Supplementary Table 1 summarises the dataset for this paper including the mean origin search output statistics. See Supplementary Tables for a description of the meaning of each variable/column. The raw input data was compiled from the Allen Ancient DNA Resource (AADR) v50 [Dav21] and modified with convertf [PPR06] and software tools from the genotype data management system Poseidon (https://github.com/poseidon-framework). We only included ancient DNA samples, and removed samples without spatial or temporal position information as well as samples outside of the defined research area (Fig. 2) and time window (median age within 8000calBC – 2000calAD).

The dataset includes both samples whose DNA libraries have undergone in-solution enrichment capture as well as samples who have been sequenced evenly across the entire genome using the so-called shotgun approach. Each sample covers an individual subset of the 1240K SNP array [Mat+15]. For quality filtering we only kept samples with 25000 or more recovered autosomal SNPs on this array, determinable molecular sex and – for male individuals – an X-chromosome contamination value (determined with ANGSD [KAN14]) smaller than 0.1. We also excluded samples that were explicitly marked as contaminated by the respective authors or assessed negatively in the AADR. In a final data filtering step we calculated pairwise distances (1 - proportion of alleles identical by state) among all samples and only kept the best preserved one from pairs/groups with distance values smaller 0.245, to remove closely related individuals or samples from the same individual.

All radiocarbon dates in the archaeological context data were recalibrated with the R package Bchron [HP08] (intercept calibration with IntCal20). Multiple radiocarbon dates for one sample were merged with sum calibration.

### Multidimensional scaling

Multidimensional scaling is a dimensionality reduction method that can be applied to genetic data to derive positions in a genetic-distance space for individual samples. Before running it on our dataset with plink --mdsplot v.1.9 [Pur+07] we removed SNPs in previously identified regions of high linkage disequilibrium within the 1240K SNP panel range according to Price et al. 2008 and Anderson et al. 2010 [Pri+08; And+10].

### Gaussian process regression

Gaussian process regression is an interpolation method for n-dimensional space. The term *Gaussian process* means that a set of observations is modelled as the outcome of a multivariate normal distribution. Each observation can be described by its mean and a covariance matrix that encodes the relations among observations. The method allows to make predictions for a dependent variable based on the position in independent variable space [Gra20]. It is a long-established method of geostatistics, where it is known as *kriging* [Mat63]. Here we treat the position in spatial space (coordinates projected to EPSG:3035) and temporal space (years calBC/calAD randomly sampled from the post-calibration radiocarbon age or uniform archaeological context age probability distribution) as three independent variables that are used to predict the dependent position on each of two multidimensional scaling result dimensions. The prediction runs for a position grid that covers the land part of the research area (resolution: 100km, 50y) with some manually set, spatial masking of low data density regions (black in Figure 5).

A crucial step in the application of Gaussian process regression is the generation of a general covariance matrix with a sensible covariance function (*kernel*) that effectively describes the degree and range of long-distance effect the model assumes for individual observations. We followed the default choice for an *anisotropic Gaussian* kernel implemented in the R package laGP v.1.5-5 [Gra16]. laGP provides comparatively fast and accurate local approximate Gaussian process modelling [Hea+19]. The default laGP kernel has the form

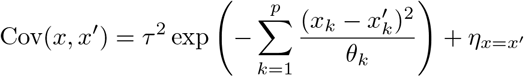

with (*x*_*k*_ − *x′*_*k*_) as the distance between all observations *x* and *x′* in the different dimensions *k* and the kernel size scaling factor *θ*_*k*_ for each dimension. *η*_*x*=*x′*_*I* is the so called *nugget* term to account for different observations of the dependent variable at the same position in independent variable space.

The two values of *θ*_*k*_ (spatial and temporal) and the value of *η* have to be fixed for the model, which is the second important decision necessary to define the covariance matrix. We applied multiple approaches (variogram analysis, maximum likelihood estimation, cross-validation) to estimate these parameters and present our results in Supplementary Text 1.

### Mobility estimation algorithm and parameter exploration

For the mobility estimation or “origin search” performed with the interpolation grid we came up with a dedicated algorithm. For each individual in the selected analysis regions we searched for the genetically closest spatial position in a timeslice of the interpolation grid *≈*700 years before their sampled age. Genetically closest means here: Has the least Euclidean distance in two dimensional MDS space. The “retrospection or rearview distance” was fixed based on the point of Cov(*x, x*^*′*^) = 0.5 on the temporal dimension. See Supplementary Text 2 for more details.

To explore the effect of 1. an MDS with three output dimensions and 2. different settings for the retrospection distance on the resulting regional mobility timeseries, we reran the analysis multiple times. See Supplementary Text 3 and the Supplementary Figures 2 and 3 for the results with these modifications.

## Acknowledgments

This research was financed by the International Max Planck Research School for the Science of Human History (IMPRS-SHH) and carried out on computational facilities of the Max Planck Institutes for the Science of Human History (MPI-SHH) and for Evolutionary Anthropology (MPI-EVA). Data collection was significantly simplified thanks to the Allen Ancient DNA Resource and the Poseidon genotype data initiative. We gratefully acknowledge insightful discussions with Joscha Gretzinger and helpful advice from Thiseas C. Lamnidis, James A. Fellows Yates, He Yu, Ayshin Ghalichi, Ke Wang (all MPI-EVA), Martin Hinz (University Bern), Martin J. Kümmel (University Jena), Oliver Nakoinz (University Kiel) and all members of the Population genetics working group at the MPI-EVA.

