## Supplementary Materials for "Estimating human mobility in Holocene Western Eurasia with large-scale ancient genomic data"

### Supplementary Figures

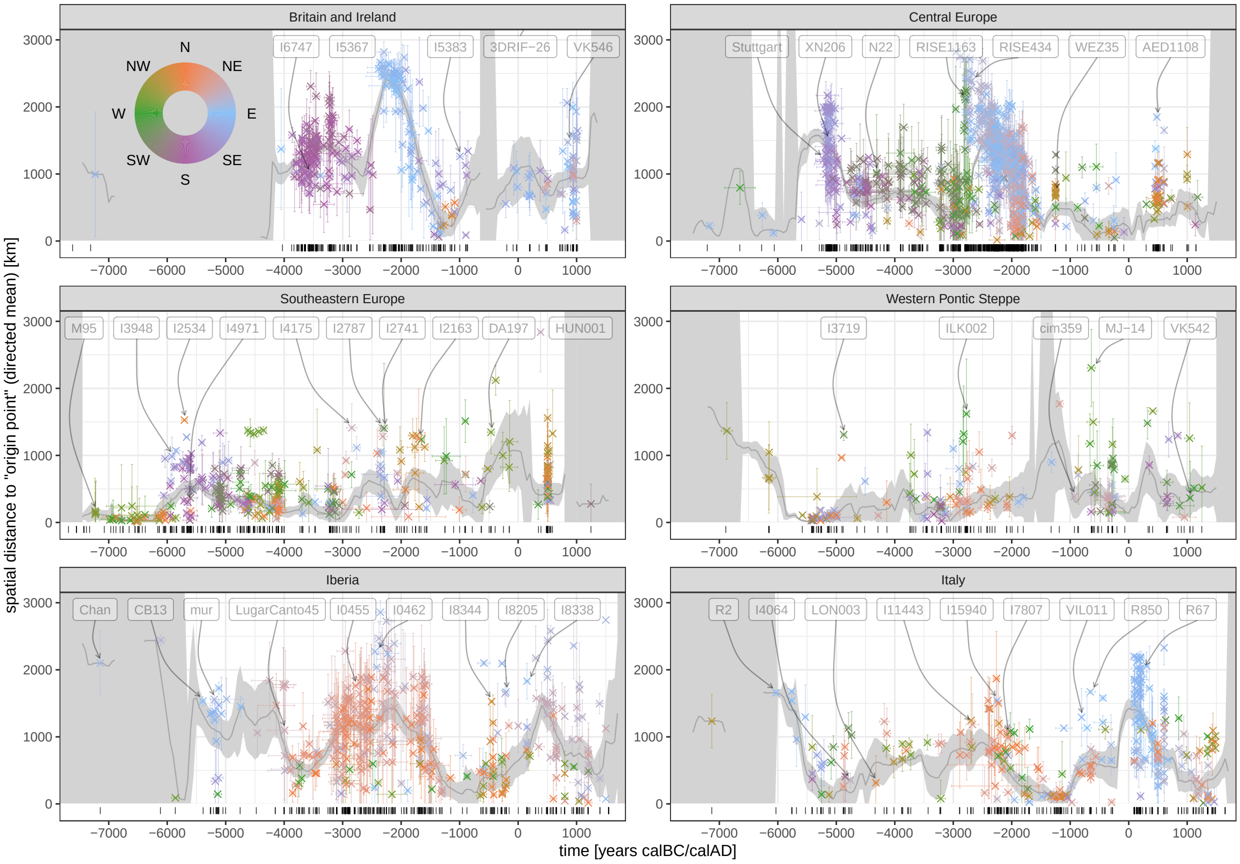


Supplementary Figure 1: Regional mobility curves for the origin search run with **three MDS output dimensions**. Beyond that as Figure 6. See Supp. Text 3 for more details.

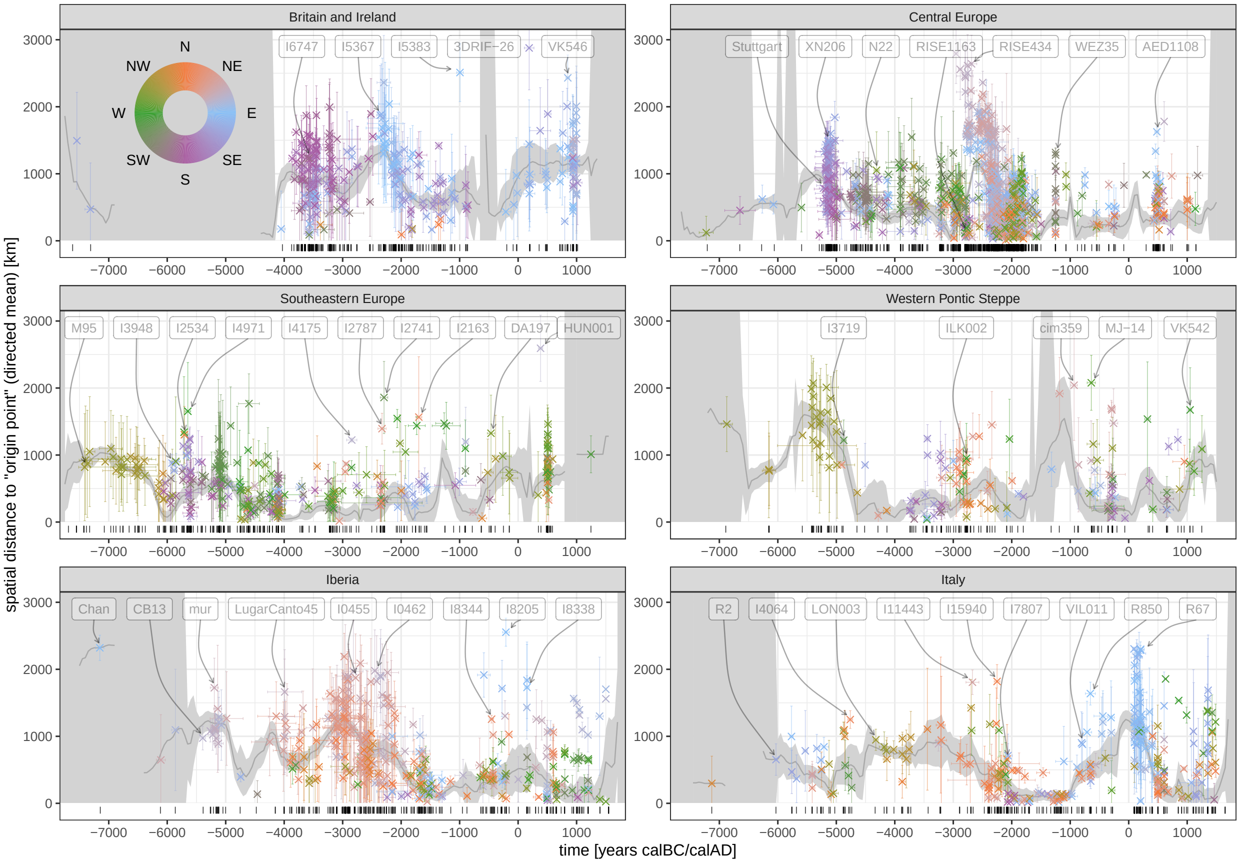


Supplementary Figure 2: Regional mobility curves for the origin search run with with a **lower retrospection distance**. Beyond that as Figure 6. See Supp. Text 3 for more details.

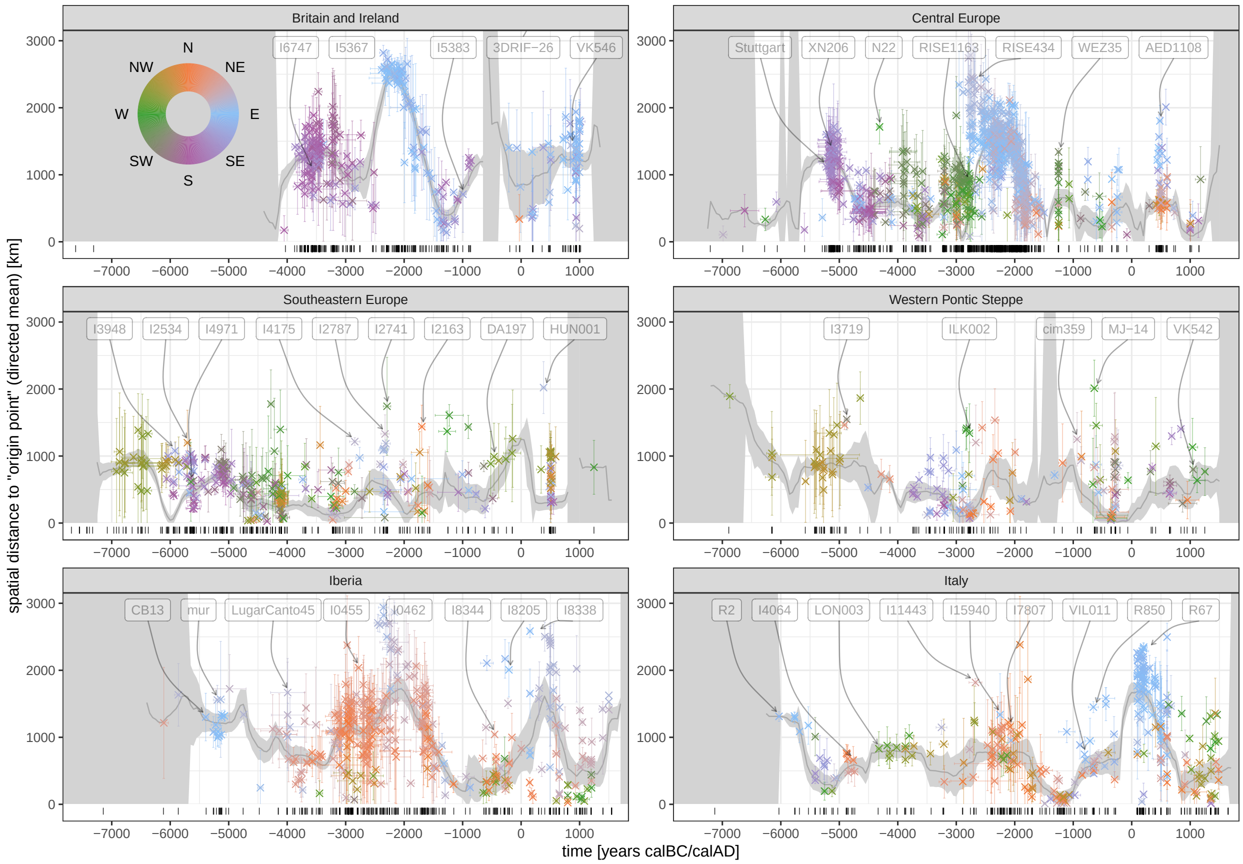


Supplementary Figure 3: Regional mobility curves for the origin search run with with a **higher retrospection distance**. Beyond that as Figure 6. See Supp. Text 3 for more details.

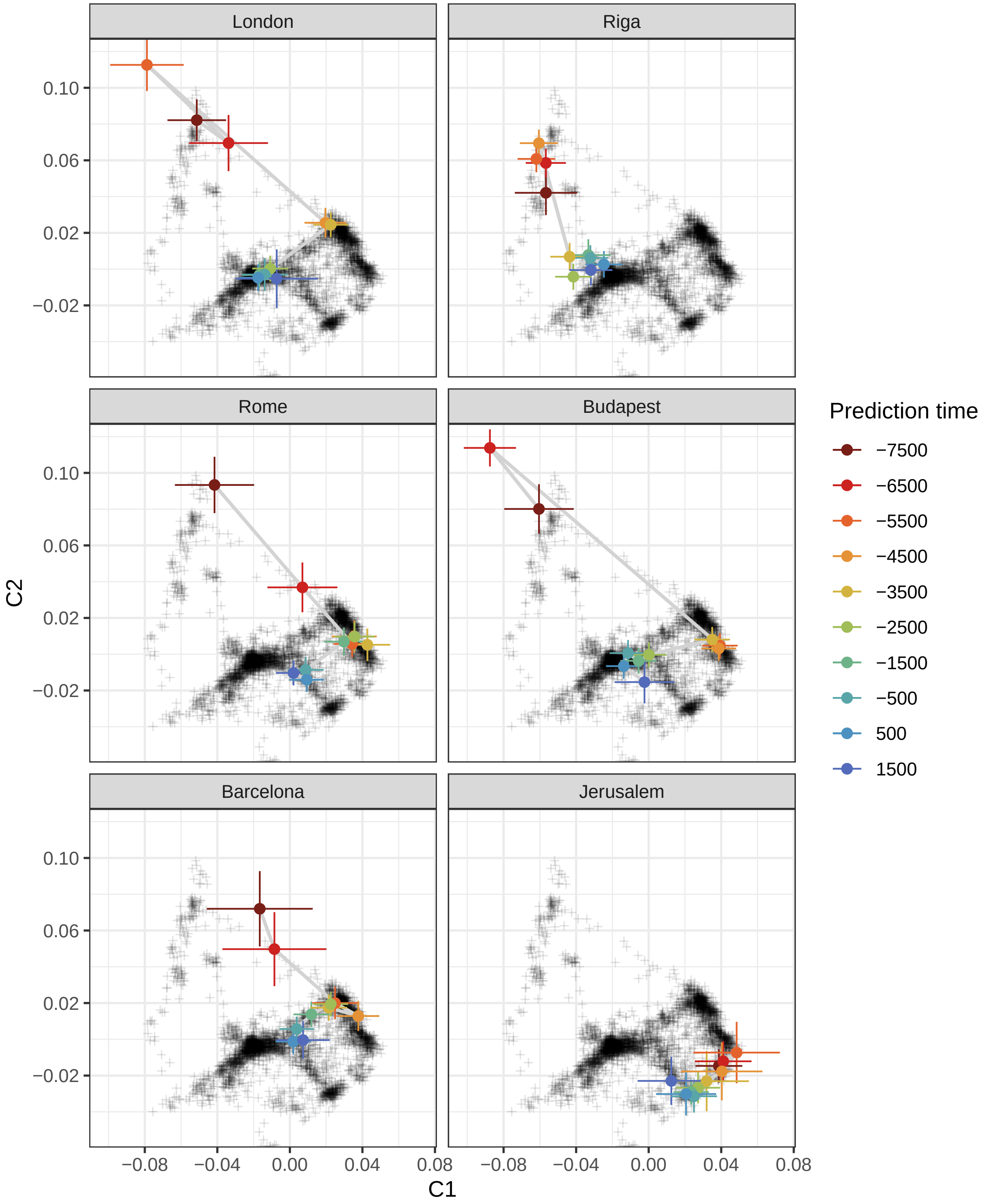


Supplementary Figure 4: Gaussian process regression based reconstruction of the past ancestry development at the spatial position of modern day city centres. Each "time-path" in multidimensional scaling space (see Figure 3) connects the interpolated positions in steps of 1000 years. The individual steps are colour-coded by age and horizontal and vertical error bars indicate the standard deviations given by the GPR model for this position. The black, semitransparent crosses in the background are the ancient samples as in Figure 3.

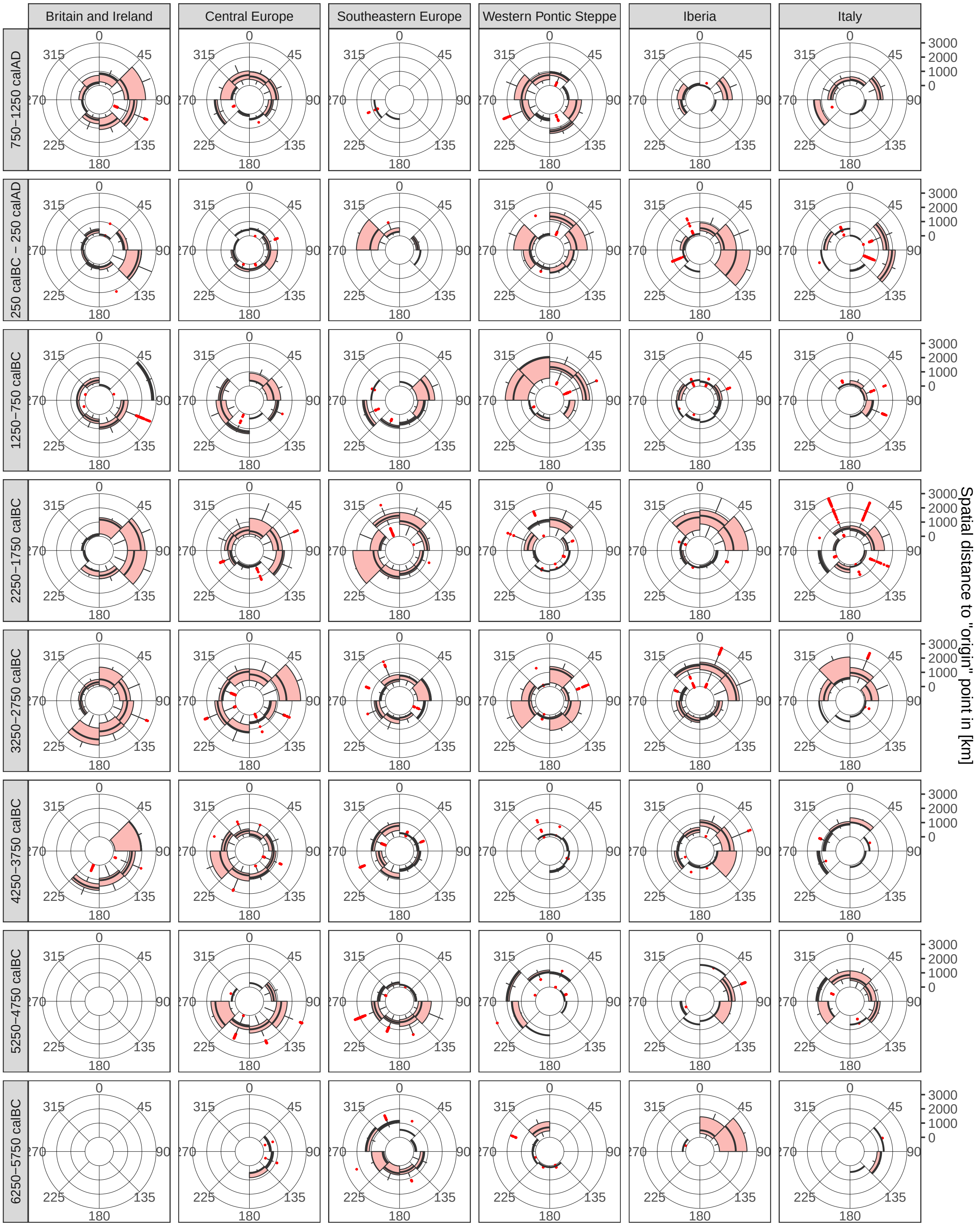


Supplementary Figure 5: Another view on the data in Figure 6. Each origin vector (100 for each input sample by temporal resampling) was attributed to a group given by the analysis region, a 500-year time window and the 45° angle range it falls in. For each of the region-time window groups there is one plot in the plot matrix. Here the distribution of directed distances within each 45° window is visualizes as a boxplot within a polar coordinate system (windrose plot). Outliers are coloured in red. The plot matrix maps time from the bottom to the top and the analysis regions on the horizontal axis from left to right.

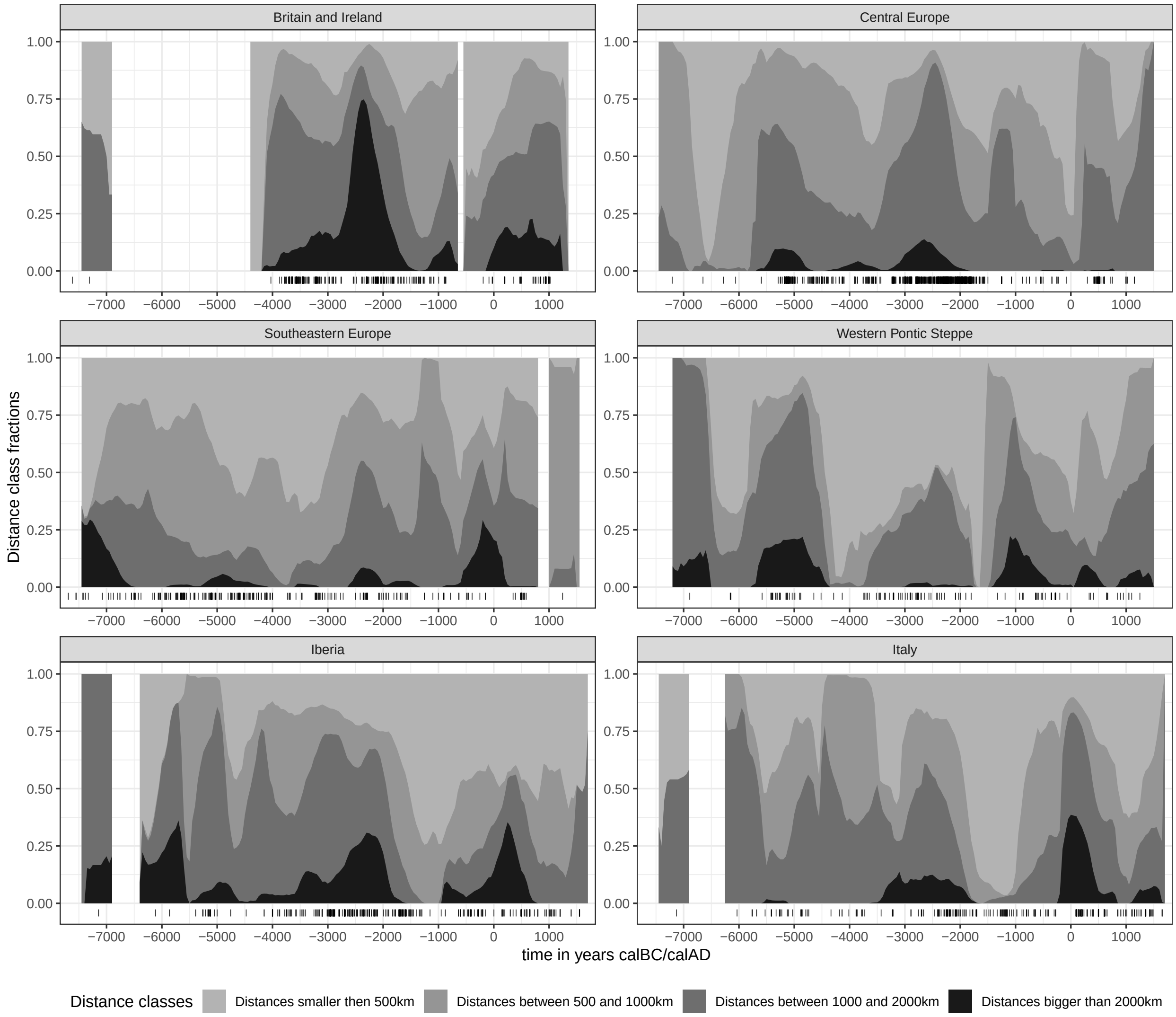


Supplementary Figure 6: Another view on the data in Figure 6. The same sliding window used to calculate the moving mean and standard deviation for Figure 6 was employed here to determine proportions of distance vectors smaller, in between and bigger 500, 1000 and 2000 kilometres. These fractions are displayed as region-wise stacked area charts. Each of the 100 temporal resampling iterations for each individual is counted separately. No-data windows are left blank.

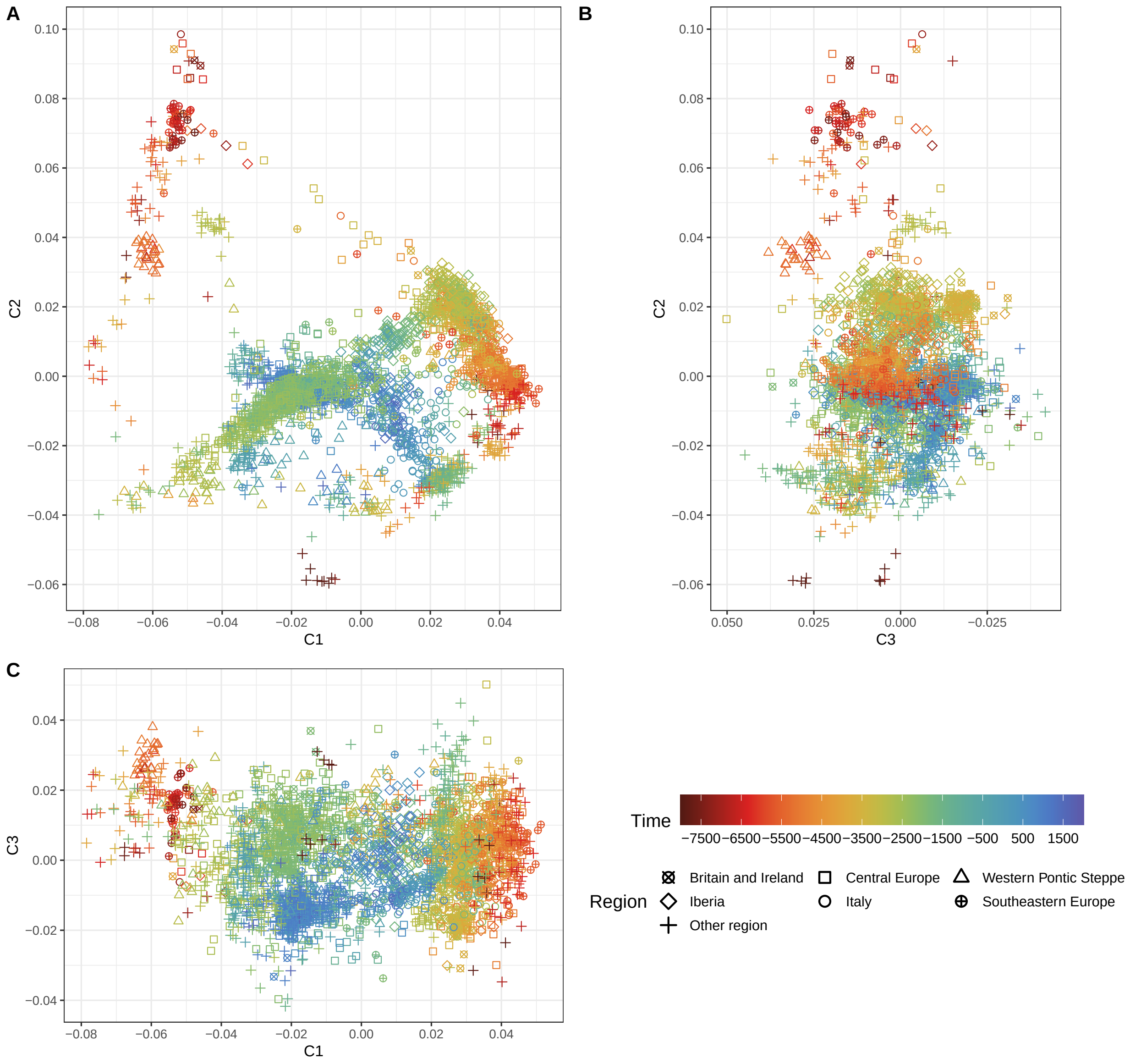


Supplementary Figure 7: Three scatter plots to show the sample distribution on the first three multidimensional scaling dimensions (see Figure 3 for the 2D version). To stay true to a 3D perspective, the printing order of each sample dot is according to the third dimension (the one not on the two axis) – with lower values always printed first. For **A** that means for example that the dots are printed in the order of their coordinate value on C3: Samples with lower values on C3 are printed first, so they are below samples with higher C3 values.

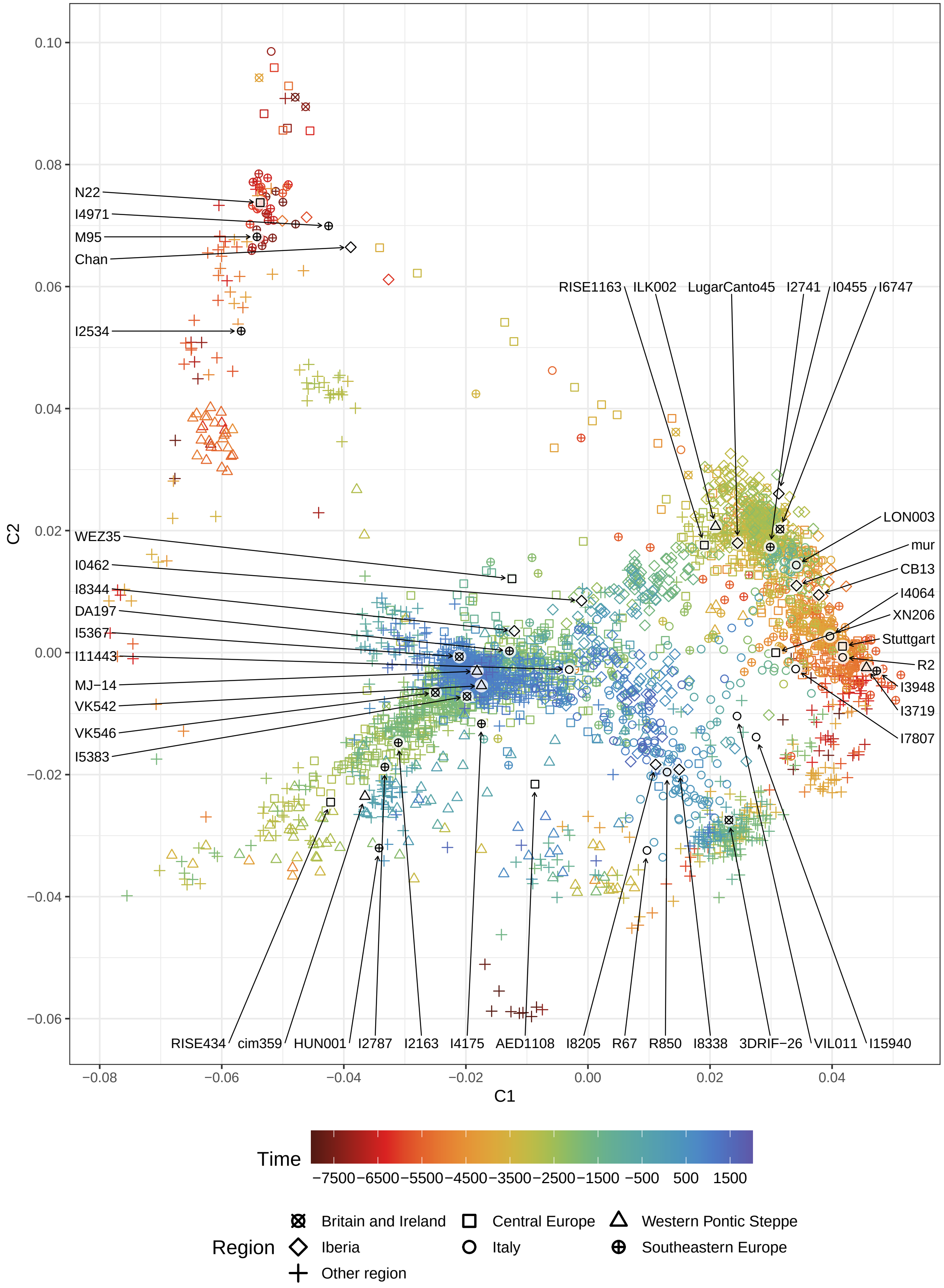


Supplementary Figure 8: A bigger version of Figure 3, where the individuals mentioned in the text (and in Figure 6) are highlighted.

### 1 Supplementary Text 1: Gaussian process regression covariance matrix parameter estimation

We consider a number of individuals distributed in space and time, with a single-dimensional (scalar) genetic PCA or MDS component as dependent variable. We use the notation $(x_i, y_i, t_i, g_i)$ to denote for each data point $i$ the set of spatial coordinates ($x_i$ and $y_i$), an age $t_i$ and the value of the genetic component $g_i$.

We intend to model our data points as a random Gaussian process, for which we are using the laGP R package for local approximate Gaussian process regression [2]. As a technical note, one of the assumptions in this package is a mean of zero in the Gaussian process, which we approximately achieve by first fitting a linear model $g \sim x + y + t$ to the data, and then considering the *residuals* instead of the original genetic values. For simplicity, and because this is a one-time operation, we just continue using the notation $g_i$, now actually denoting the residuals of the linear model instead of the raw genetic component.

A key ingredient for Gaussian process regression is the covariance kernel function, for which we here follow the standard choice of a squared exponential, which in general terms for p-dimensional input data and in the notation of laGP is defined as

$$Cov(x, x') = \tau^2 \exp\left(-\sum_{k=1}^{p} \frac{(x_k - x'_k)^2}{\theta_k}\right) + \eta\delta(x - x') \tag{1}$$

where $x$ and $x'$ are positions in p-dimensional space, and $\sqrt{\theta_k}$ are lengthscale parameters for each dimension. $\eta$ is an additional noise term to be added only if $x = x'$, using the delta-distribution, the so called nugget term. $\tau^2$ is a general scaling parameter.

Specifically for the purpose of spatio-temporal modelling with isotropic space, we write this covariance function as

$$\mathrm{Cov}(r, u) = \tau^2 \exp\left(-\left(\frac{r}{\rho}\right)^2 - \left(\frac{u}{\alpha}\right)^2\right) + \eta\delta(x - x') \tag{2}$$

where we have changed notation slightly, and introduced the spatial kernel radius $\rho$ and the temporal kernel radius $\alpha$, which by construction have now length- and time-dimensions (measured in years and kilometres, respectively). For readers interested in mapping notation to the laGP documentation, we note that in their notation the correlation function $K(r, u)$ is defined as the normalized part of the covariance, such that $\mathrm{Cov}(r, u) = \tau^2 K(r, u)$.

#### 1.1 Variogram analysis

One possibility to inspect plausible parameters for $\tau$, $\eta$, $\rho$ and $\alpha$ as defined in 2 is variogram analysis (see also [3], [1]).

It is instructive to first consider variograms in the context of continuous fields, where the field value $g(x, t)$ is defined at all spatial points $x$ (which in our case are two-dimensional) and all time points $t$.

The semivariogram is then defined as the mean squared difference of field values at given spatial and temporal distances:

$$V(r, u) = \frac{1}{2}\langle (g(s, t) - g(s + r, t + u))^2 \rangle_{s,t} \tag{3}$$

where the average runs over all space-time points $(s, t)$.

Under the assumption of constant variance $\langle g(s,t)^2 \rangle = \langle g(s+r, t+u)^2 \rangle$ for all $s, r, t, u$, we can establish the relationship of the semivariogram and the covariance function of the Gaussian process:

$$\begin{aligned} V(r,u) &= \frac{1}{2}\langle (g(s,t) - g(s+r, t+u))^2 \rangle_{s,t} \\ &= \frac{1}{2}\langle g(s,t)^2 - 2g(s,t)g(s+r,t+u) + g(s+r,t+u)^2 \rangle \\ &= \frac{1}{2}\langle g(s,t)^2 \rangle - 2\langle g(s,t)g(s+r,t+u) \rangle + \langle g(s+r,t+u)^2 \rangle \\ &= \mathrm{Cov}(0) - \mathrm{Cov}(r,u) \end{aligned} \tag{4}$$

with $\mathrm{Cov}(r,u) = \langle g(x,t)g(x+r,t+u) \rangle$

So the variogram is directly related to the covariance of the Gaussian process:

$$V(r,u) = \mathrm{Cov}(0) - \mathrm{Cov}(r,u) \tag{5}$$

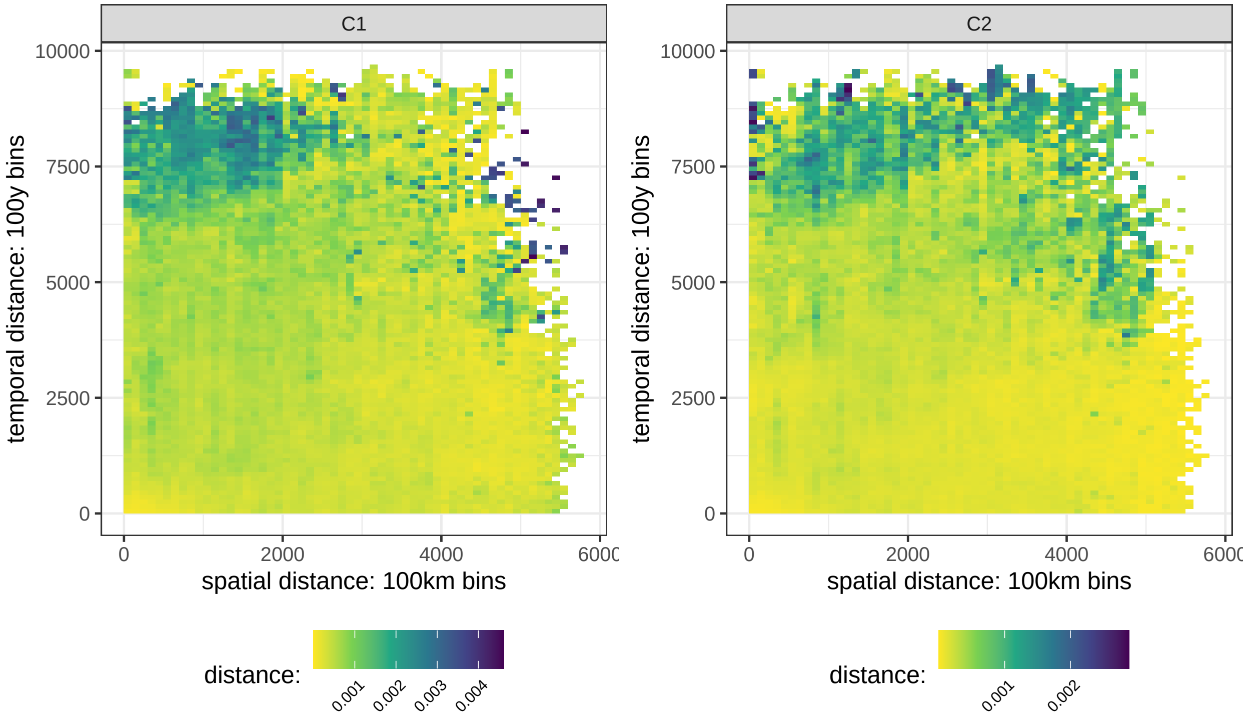


Supp. Figure I: Empirical semivariogram rasters calculated including all samples in the analysis dataset, with one plot for each ancestry component (MDS coordinate) C1 and C2. The fill colour represents the mean squared pairwise distance in the respective space-time bin. For some basic detrending these distances were calculated not directly on the ancestry components, but on the residuals of a simple linear model, where the genetic coordinates for each sample are predicted by their spatiotemporal position.

Following [1] p.30, the **empirical semivariogram**, defined for a set of actual datapoints, can be computed as a binned version of the continuous semi-variogram definition employed above. Specifically, instead of continuous spatial and temporal "radius" values $r$ and $u$, as in the continuous version, we now consider bins $R_k = (r_k, r_{k+1})$ and $U_l = (u_l, u_{l+1})$, with boundaries $r_1 < r_2 < \ldots$ and $u_1 < u_2 < \ldots$. We then write

$$V(k,l) = \frac{1}{2N(k,l)} \sum_{i,j} (g_i - g_j)^2 I\left(\sqrt{(x_i - x_j)^2 + (y_i - y_j)^2} \in R_k, |t_i - t_j| \in U_l\right) \tag{6}$$

where $I(\text{condition})$ is an indicator function that is 1 if the condition is true and zero otherwise, and the normalization $N$ is

$$N(k,l) = \sum_{i,j} I(\sqrt{(x_i - x_j)^2 + (y_i - y_j)^2} \in R_k, |t_i - t_j| \in U_l) \tag{7}$$

Supp. Figure I is one way to visualize $V(k,l)$ as a raster plot. The bins $R_k$ and $U_l$ are here chosen such that $r_i - r_{i-1} = 100\text{km}$ and $u_l - u_{l-1} = 100\text{years}$.

Another way to visualize relevant information from this empirical semivariogram is to select pairwise distances from within small stripes along the x- and y-axis in Supp. Figure I as we did for Supp. Figure II. **A** contains all genetic distances for very low ($< 50$y) temporal distance, **B** for very low spatial distance ($< 50$km).

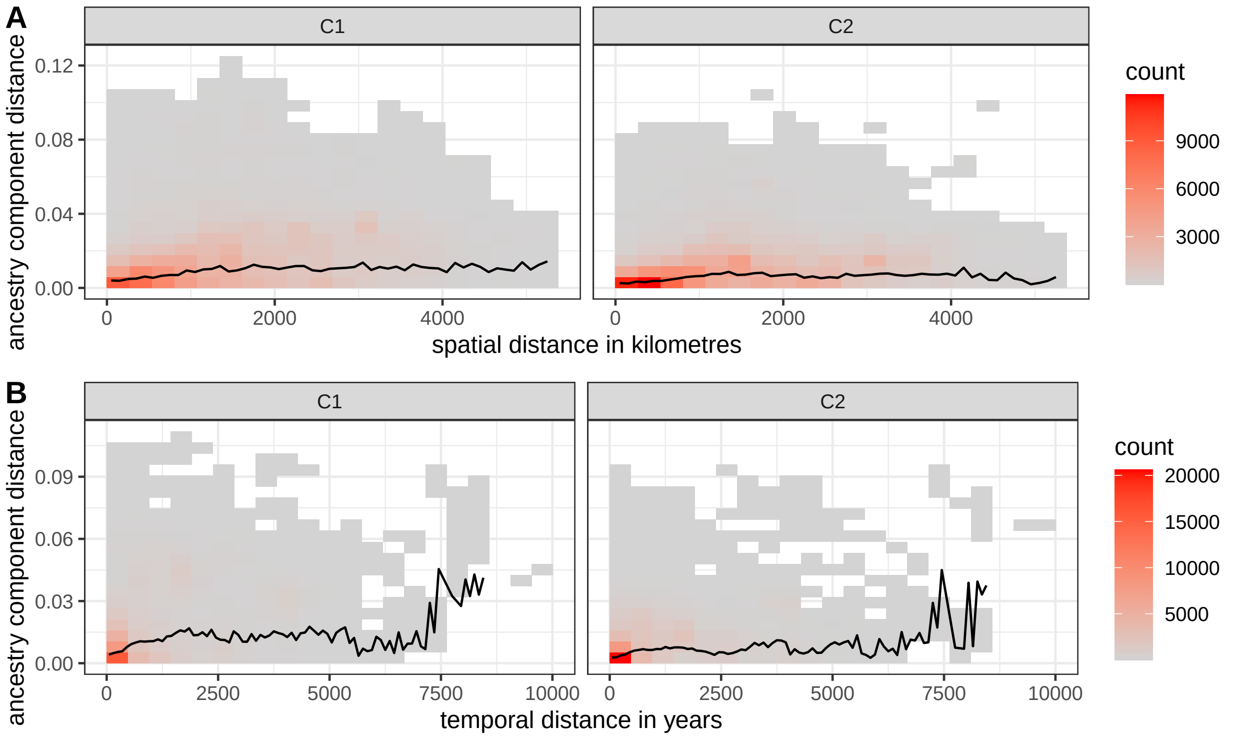


Supp. Figure II: Another view on the semivariogram: Counts of all pairwise distances along the ancestry components for very low temporal (**A**) or spatial (**B**) distances ($< 50$years $\|$ $< 50$km).

To finally determine the kernel parameters ($\rho$ and $\alpha$ or $\theta$) from the this empirical semivariogram on residuals we attempted to fit a squared exponential function to it (as in equation 2). Unfortunately we came to the conclusion that we cannot do so unambiguously. We cannot co-estimate Cov(0) and the kernel radiuses simultaneously. Supp. Figure III illustrates this. It shows only a single cut through the semivariogram, and two vastly different exponential models that look almost the same.

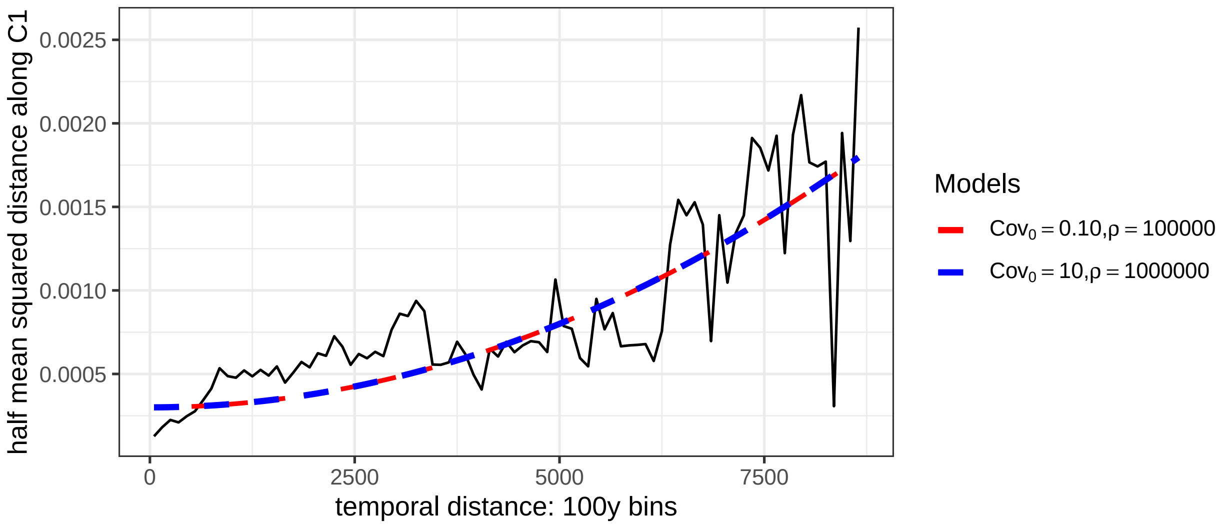


Supp. Figure III: A variogram for one timeslice ($x \in [0, 100]$) with two different, but equally well fitting exponential models.

Indeed: For $(r/\rho) \ll 1$ we have

$$\begin{aligned} \mathrm{Cov}_0 - \mathrm{Cov}_0 * \exp(-(r/\rho)^2) &= \\ \mathrm{Cov}_0 \left(1 - \exp(-(r/\rho)^2)\right) &\approx \\ \mathrm{Cov}_0 \left(1 - (1 - (r/\rho)^2)\right) &= \\ \frac{\mathrm{Cov}_0}{\rho^2} r^2 & \end{aligned} \tag{8}$$

where we have used the Taylor expansion for the exponential function: $\exp(x) = 1 + x + \mathcal{O}(x^2)$. So for small values of $r/\rho$ we get an approximate squared function with a coefficient of $\mathrm{Cov}_0/(\rho^2)$, which shows that the model is approximately invariant under changes of $\mathrm{Cov}_0$ and $\rho^2$ that leave the ratio constant. This is the case in the two curves above. We don't get into the plateau of the variogram, so can not fit $\mathrm{Cov}_0$ and $\rho$ independently. We concluded that empirical variograms can not be used for kernel length estimation in this particular context, and turned to different estimation approaches below.

However, the variogram at least exposes an approach to estimate the variance $\tau^2$ and nugget term $\eta$ (as in equation 1). First, from the form of the covariance function 2 we have $\mathrm{Cov}(0,0) = \tau^2(1+\eta)$. At the same time, for small but non-zero values of $r$ and $u$ we have $\mathrm{Cov}(r \to 0, u \to 0) = \tau^2$. So for the semivariogram we get:

$$V(r \to 0, u \to 0) = \mathrm{Cov}(0,0) - \mathrm{Cov}(r \to 0, u \to 0) = \tau^2(1+\eta) - \tau^2 = \tau^2\eta \tag{9}$$

so for the nugget term we have now an estimator

$$\hat{\eta} = \frac{V(r \to 0, u \to 0)}{\tau^2} \tag{10}$$

This can be readily derived, since the variance $\tau^2$ can be estimated as the overall sample variance of the data, i.e.

$$\hat{\tau^2} = \frac{1}{N}\sum_i (g_i - \bar{g})^2 \tag{11}$$

where $N$ is the number of data points and $\bar{g}$ is the mean genetic value.

Supp. Figure IV shows the distribution of pairwise squared genetic distances for samples that are less then 50 years and 50 kilometres apart. Each distance value is scaled according to the estimator defined in equation 10 and the mean of these pairwise distances is a good default for the nugget term. We therefore conclude that a nugget of $\eta = 0.07$ is suitable for both ancestry components, and have fixed this value for most subsequent analyses.

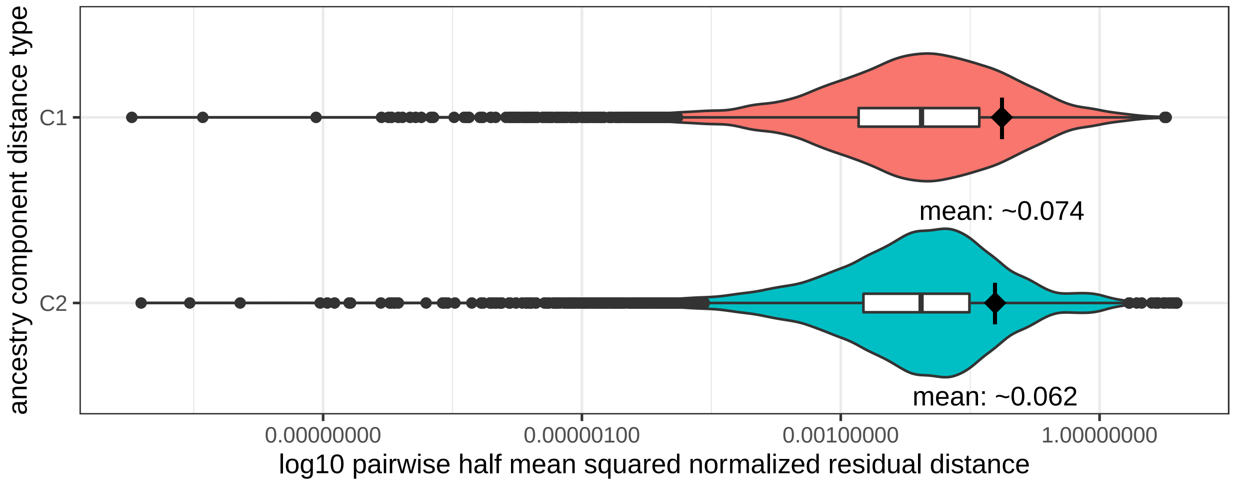


Supp. Figure IV: Violin- and boxplot of the detrended pairwise distance distribution for different ancestry components in a short and narrow temporal and spatial distance window ($<$ 50km & $<$ 50years). The diamond shape is positioned at the mean point of the distribution. See Supp. Fig. XVI for C3.

#### 1.2 Maximum likelihood estimation

The laGP package [2] provides two different maximum likelihood estimation (MLE) algorithms for automatic kernel parameter exploration in anisotropic spaces: `mleGPsep` and `jmleGPsep`. According to the manual `mleGPsep` uses L-BFGS-B optimization (a limited memory quasi-Newton approximation of the Broyden-Fletcher-Goldfarb-Shanno algorithm) to get an estimate of $\theta$ ($\rho$ and $\alpha$ above). It allows for joint estimation of $\theta$ and the nugget $\eta$ with a common gradient. `jmleGPsep` on the other hand is explicitly designed for joint inference by iterating over the marginals of $\theta$ and $\eta$. laGP allows to set starting parameters and search boundaries for both algorithms with the helper functions `darg` and `garg`. According to the manual, these ”leverage crude summary statistics” over the independent and dependent input variables to define sensible defaults.

`mleGPsep` and `jmleGPsep` as implemented in laGP are generally not well suited for spatiotemporal data without inherent latitudinal or longitudinal bias, as they optimize each input dimension separately: Instead of one spatial kernel radius $\theta_s$ and one temporal kernel radius $\theta_t$, they effectively yield two separate values for $\theta_s$, one for the spatial x axis ($\theta_x$), and one for the spatial y axis ($\theta_y$). Despite this, we decided to apply the algorithms here to get a first estimate for $\theta$ and to test our previous conclusion concerning $\eta$.

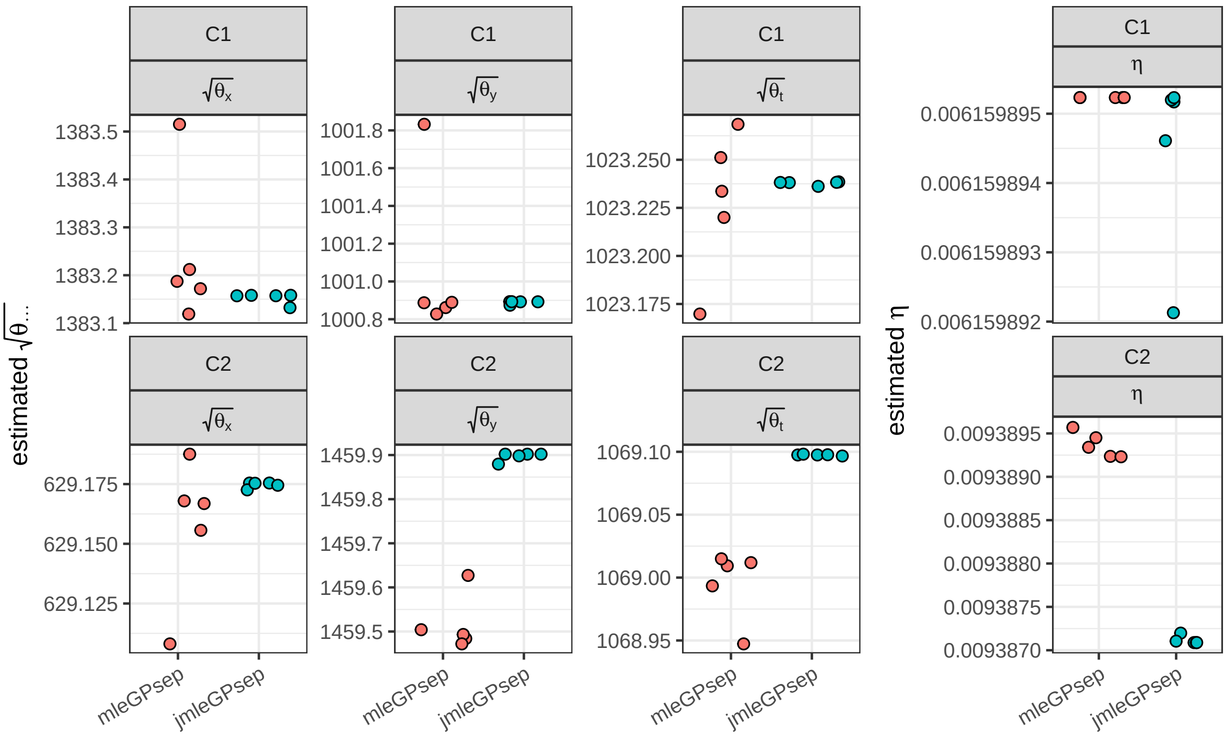


Supp. Figure V: Results of the kernel parameter estimation with the laGP maximum likelihood algorithms `mleGPsep` and `jmleGPsep`. Each dot (horizontally jittered) represents the result of one run for one parameter. For each permutation of algorithm and ancestry component 5 runs were calculated. Of the 20 total runs none had to be removed as they all converged.

Supp. Figure V shows the result of multiple runs for each combination of algorithm and ancestry component. The estimated $\theta$ values for the three dimensions are very similar between the two algorithms (`mleGPsep` and `jmleGPsep`) but different for the two ancestry components modelled with the Gaussian process (dimensions C1 and C2 in MDS space). Note that we report $\sqrt{\theta}$ instead of $\theta$, since that has the more interpretable unit (kilometres and years, respectively), see equation 1. The values are relatively large, but seem at least generally plausible, given how far the influence of each point could "radiate" in a squared exponential model and how far prehistoric interaction networks may have spanned (see Supp. Figure VI to get some intuition). One consistent observation is that $\theta_x$ should be different from $\theta_y$ for a good model. So the above mentioned anisotropy issue does indeed affect the outcome of the parameter estimation and poses a form of overfitting. We do not believe though that a model with a latitude-longitude mismatch is justified in this context.

The estimated values for $\eta$ are about one order of magnitude smaller than the ones estimated from the variogram. We assume this to be an effect of the implausible anisotropy. Experimental interpolation test runs with $\eta < 0.04$ led to overfitting in settings with fixed $\theta_x == \theta_y$ and we therefore decided to keep $\eta$ fixed as decided above.

laGP also provides the function `mleGP` to estimate $\theta$ and $\eta$ in isotropic systems and we decided to employ this algorithm as well. To account for the anisotropic nature of the space-time relationship we introduced a scaling factor that manipulates the temporal position information. Starting from the default 1 (1km = 1y) we increased and decreased the scaling factor in a rescaling sequence from 1/10 to 2. Supp. Figure VII

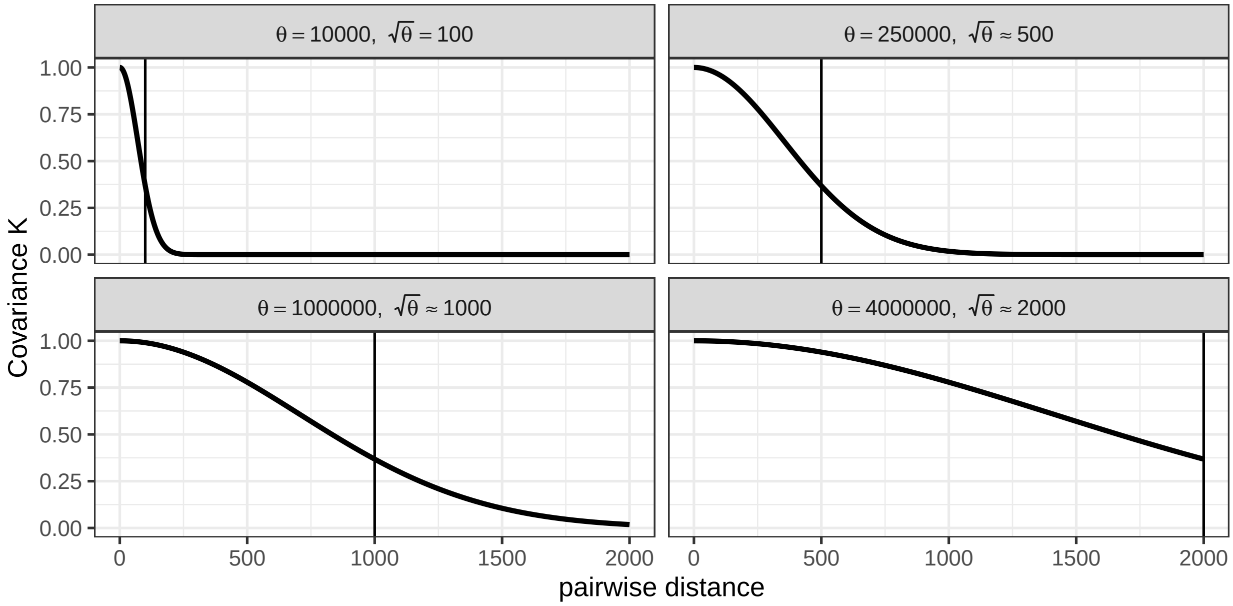


Supp. Figure VI: Example curves to illustrate the behaviour of a squared exponential function $K_{ij} = \exp(\frac{-\|x_i - x_j\|^2}{\theta})$ with different values of $\theta$. The "pairwise distance" could for example be in kilometres or years.

documents the result: `mleGP` yields only one value for $\theta$, which reacts to the forced temporal "contraction" and "inflation". One way to imagine this is a rigid sphere in a changing cuboid universe: We contract or inflate the cuboids z-axis and estimate for each setting **1.** if a sphere is a good assumption for predicting observations (Supp. Figure VII **B**) and **2.** which radius the sphere should ideally have (Supp. Figure VII **A**). As stated above, we used the fixed value for $\eta$ here.

For both ancestry components, C1 and C2, increasing and decreasing the scaling factor, so temporal inflation and deflation, quickly deteriorates the model likelihood. A scaling factor of 1, so $\theta_t = \theta_s$ and 1km = 1y, yields good results for both (only for C2 we observe a tiny increase in likelihood for smaller scaling factors). The estimates for the absolute values of $\sqrt{\theta}$ are smaller, but on the same magnitude as for the anisotropic estimation above.

#### 1.3 Crossvalidation

As a third and more independent method to estimate $\theta$, we turned towards a simple crossvalidation approach, which allows to see the effect of different kernel size values on prediction accuracy and precision. We explored a $\theta$ grid with 40 values for the spatial kernel size $\sqrt{\theta_s} = 50, 100, 150, ..., 1900, 1950, 2000$ km and 40 values for the temporal kernel size $\sqrt{\theta_t} = 50, 100, 150, ..., 1900, 1950, 2000$ years. The nugget term $\eta$ was again fixed as decided above. Our crossvalidation algorithm includes the following steps and was applied for each ancestry component and $\theta_s$ and $\theta_t$ combination separately:

1. Randomly reorder the observations

2. Split the observations into 10 groups

3. Build a laGP GPR model from 9 of the 10 groups and use it to predict the 10th. Do this for all

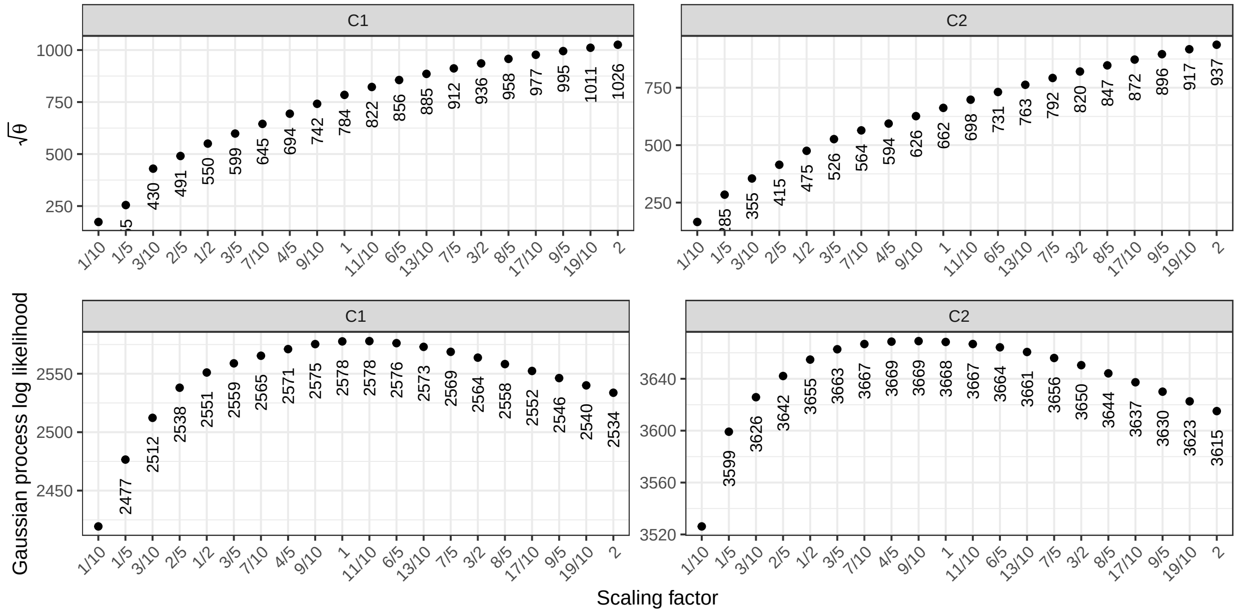


Supp. Figure VII: Results of `mleGP` exploration runs with variable scaling of temporal and geospatial space. `mleGP` assumes an isotropic sytem.

combinations of groups

4. Calculate the distance between real and predicted value for each observation

As these steps are also repeated 10 times, this crossvalidation is computationally very expensive and was calculated on a high performance computing cluster. The fast approximate GPR implementation in laGP helped substantially to make this feasible.

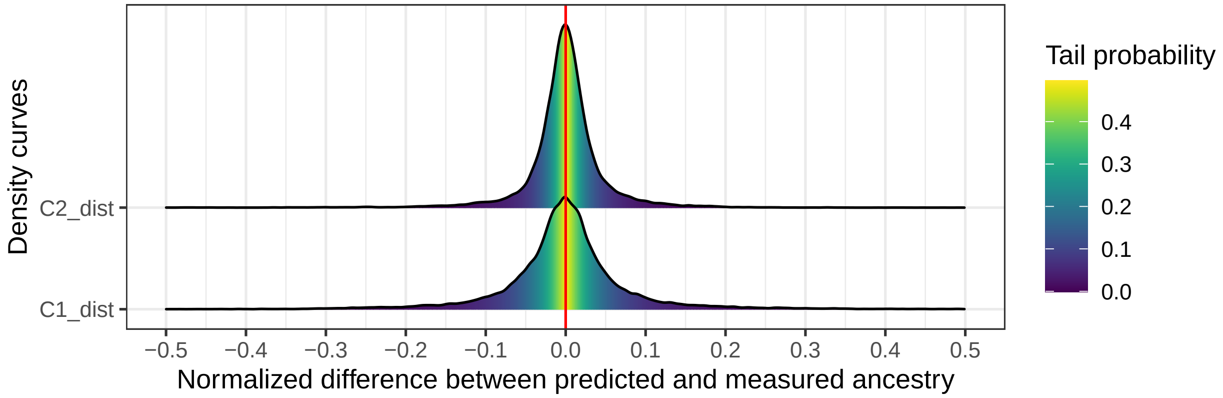


Supp. Figure VIII: Distribution of 100,000 randomly drawn deviations out of all crossvalidation prediction-observation distances. The distance values were normalized to the total range of the respective ancestry component.

Supp. Figure VIII shows the distributions of a large sample of normalized (to the range of the ancestry components) distance values. As expected, the distances form a distribution around zero. Most predictions are in a 10% margin around the observed ancestry data. That means that **1.** the GPR models are generally good at predicting the ancestry of unknown observations and **2.** there must exist multiple combinations of $\theta_s$ and $\theta_t$ that yield a solid GPR model.

The latter is confirmed when we look at the mean squared difference rasters in Supp. Figure IX **A** and **B**. Generally, good predictions are possible in a remarkably large corridor of $\theta_s$ and $\theta_t$ value permutations.

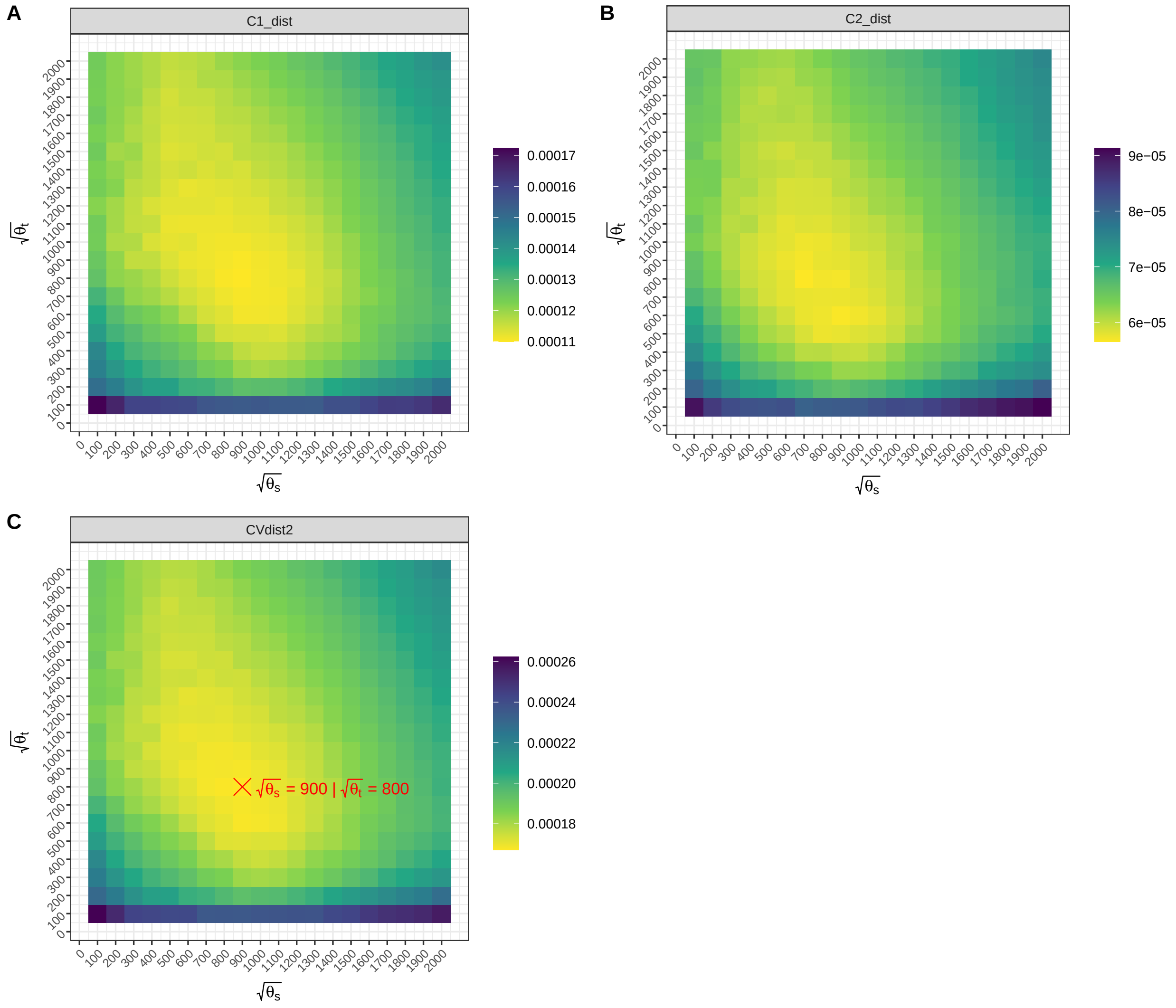


Supp. Figure IX: Crossvalidation results (mean squared differences between prediction and observation) for both ancestry components and different $\theta_s$ and $\theta_t$ combinations. Plot **C** shows a merged result for both ancestry components. The combination of $\theta_s$ and $\theta_t$ with the best mean predictive power is highlighted in red.

For Figure IX **C** we calculated the differences as Euclidean distances across both ancestry component differences in **A** and **B**. The red cross in the figure marks the lengthscale parameter combination with the best predictive capabilities for both ancestry components C1 and C2 combined.

#### 1.4 Decision

We believe that a large kernel with $\theta_t \approx \theta_s$ as indicated by maximum likelihood estimation and crossvalidation has the best mean postdictive power for the European spatiotemporal ancestry field given the current amount and distribution of data. The analysis in this paper therefore relies on the kernel settings estimated via crossvalidation. See Supp. Table I for a summary of the parameters we chose – also for the retrospection distance parameter, which will be introduced in Supp. Text 2, and the parameter choices for the experiments in Supp. Text 3.

Beyond that we also experimented with smaller kernels and kernels with $\theta_s \ll \theta_t$ and $\theta_s \gg \theta_t$. Above results indicate that a rather large range of covariance functions may yield satisfying models, and for the mobility estimation attempted here, smaller kernels may theoretically be more useful. They could produce stronger and more sharply bounded signals for specific events of change. A kernel with a high $\theta_s$ and $\theta_t$ on the other hand may obscure phenomena of temporal change by smoothing them out and by artificially attributing them an earlier starting and later end time. In the end, though, our experiments left us to believe that the different plausible kernel choices yield rather similar patterns for the mobility estimation and we focused on only one setting.

| | $\sqrt{\theta_s}$ lengthscale space [km] | $\sqrt{\theta_t}$ lengthscale time [years] | $\eta$ nugget | $u$ retrospection distance [years] |
|---|---|---|---|---|
| Default | 900 | 800 | 0.07 | 667 |
| 3D MDS | 400 | 1100 | 0.22 | 916 |
| High retrospection distance | 900 | 800 | 0.07 | 942 |
| Low retrospection distance | 900 | 800 | 0.07 | 430 |

Supp. Table I: Overview of the main parameters chosen for the analysis in the main text (Default) and the experiments in Supp. Text 3.

### 2 Supplementary Text 2: The mobility estimation algorithm

The main question for this paper was to estimate and quantify human mobility through time and space from genetic data. We assume this can in principle be done because people carry their genetic ancestry profiles with them when they move. Mobility estimation requires **1.** a suitable dimension reduction for "ancestry", and **2.** a handle on the sparseness of genetic data through space and time.

We deal with the first requirement by multidimensional scaling, which assigns every individual two or more principal components. For simplicity, in the following we will just assume a single principal component, called $C$. In our implemented application we use two or three components and calculate distances in genetic space as 2D or 3D Euclidean distances.

The second requirement can be solved by interpolation through Gaussian Process Regression, as implemented in the laGP R package [2]. With a suitable kernel as determined in Supplementary Text 1, this yields an estimate of the genetic ancestry component $C$ as a *smooth* function in space and time. "smooth" here means that our function $C$ is continuous and differentiable within the focus area and focus time.

#### 2.1 Ancestry fields

Consider now this genetic component $C$ as a function of a time variable $t$ and a single space dimension $x$. Then, our genetic component is a function in space and time, i.e. $C(x, t)$, like for example a temperature "field". At a particular time point, it might look like Supp. Figure X.

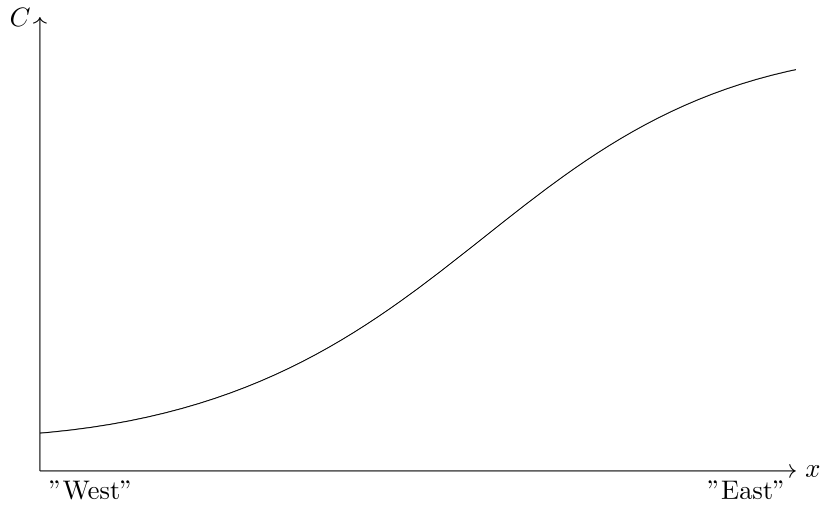


Supp. Figure X: Schematic of the spatial distribution of a genetic ancestry component $C$ at a given point in time.

In this example, the genetic component $C$ follows a gradient with lower values on the left side (say, "West") and higher values on the right side (say, "East").

#### 2.2 Sample-wise mobility

Now, let us consider an archaeogenetic sample $S_{x_1,t}$, so a measurement of the ancestry $C$ at a given point in space and time (see Supp. Figure XI). $S_{x_1,t}$ is theoretically expected to be on the $C(x, t)$-curve. For real-world data it is usually not, though. Instead the ancestry profile measured in $S_{x,t}$ may deviate from the one postdicted by the ancestry field. But the ancestry profile in $S_{x_1,t}$ could exist at this time somewhere on $C(x,t)$ at another point in space. The spatial distance $\Delta x$ between $S_{x_1,t}$ and $C(x,t)$ therefore becomes a measure for the deviation of expected and measured ancestry.

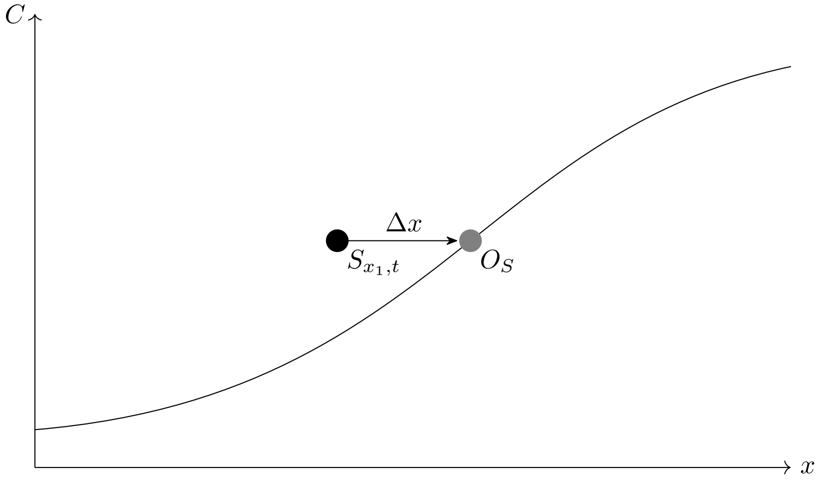


Supp. Figure XI: A sample S is added to the previous drawing. It does not fall on the curve, but its ancestry profile exists there in some spatial distance $\Delta x$.

We can go one step further and calculate this origin distance for $C(x,t-u)$, so for $C$ at a time $t-u$ before the individual behind $S_{x_1,t}$ lived. $\Delta x$ can then be considered a proxy for mobility involving or surrounding the individual represented by $S$. If $\Delta x \approx 0$, then no spatial mobility took place within the time $u$. For $\Delta x \gg 0$ we can assume some relocation. $u$ is a free parameter and we call it the retrospection or rearview distance. The vector with length $\Delta x$ we call the mobility- or origin vector $\overrightarrow{x_S}$. It has both an informative length/magnitude and direction.

Supp. Figure XII depicts the case that the exact ancestry profile of $S_{x_1,t}$ does not exist on $C(x,t-u)$. In our implementation we then calculated $\overrightarrow{x_S}$ to the most closely related ancestry profile.

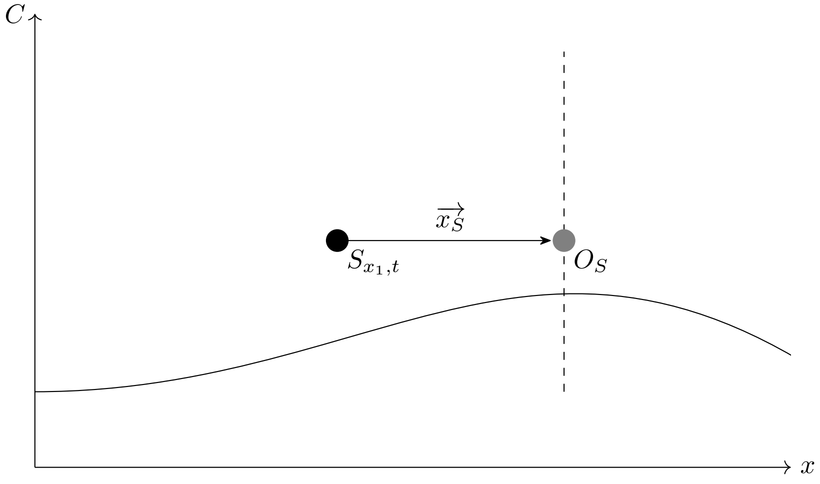


Supp. Figure XII: Here the exact ancestry profile of $S$ is not available on the $C$ curve. But there is a point where the difference is minimal.

This case is indeed the normal mode of operation, as the ancestry field is in fact represented by a discrete grid of spatiotemporal positions. Supp. Figure XIII gives an idea how the search algorithm operates for each sample:

1. The sample $S$ is attributed a previous time slice $t - u$ in the ancestry field grid.
2. Within this grid slice the spatial position with the least "genetic" distance (so euclidean distance in MDS space) is determined.
3. $\overrightarrow{x_S}$ is calculated and stored.

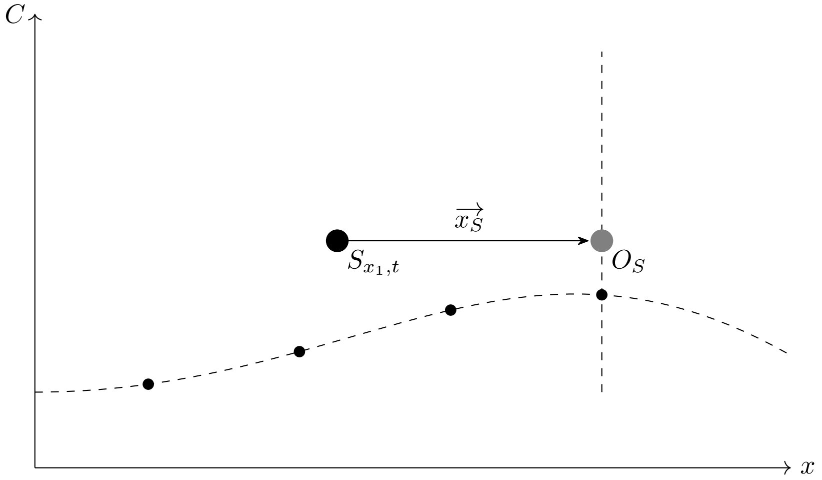


Supp. Figure XIII: $C$ is not constructed as a continious function in our implementation, but as a discrete grid of interpolated field values.

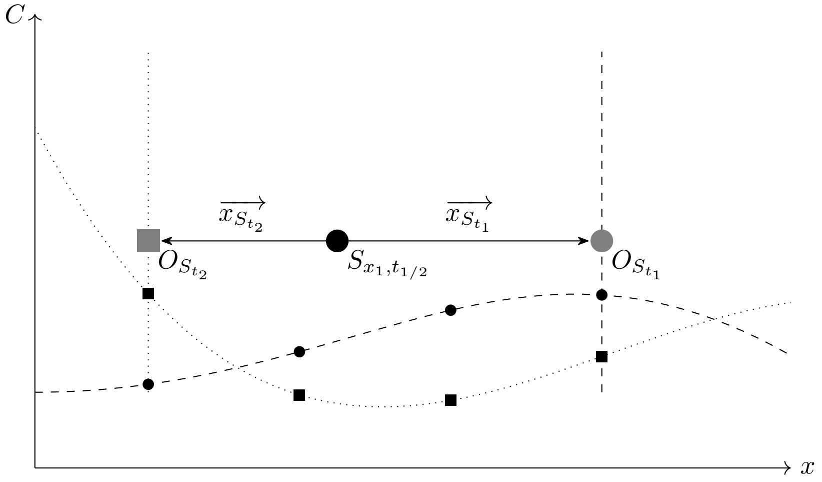


Supp. Figure XIV: Multiple $\Delta x$ values are determined for each $S$, as the temporal coordinates of all input data points are resampled multiple times. That causes different interpolations of $C$ (here: the dotted and the dashed curve), and thus different origin points $O$.

To account for dating uncertainties, we ran the ancestry search for a given sample not only once, but multiple times, where the age of all input observations for the field generation was resampled from the (post-calibration) dating probability distributions. That yields multiple vectors $\overrightarrow{x_{S_{t_1}}}, ..., \overrightarrow{x_{S_{t_n}}}$ for each input sample $S$ (see Supp. Figure XIV). We consider this a "natural" way to obtain a distribution of $\Delta x$ vectors, indicating the precision of a given, individual-wise mobility estimate.

#### 2.3 Mobility proxy

As we have two spatial dimensions to consider in the real-world data, $\overrightarrow{x_S}$ is actually a two dimensional vector. From each distribution $\overrightarrow{x_{S_{t_1}}}, ..., \overrightarrow{x_{S_{t_n}}}$ we can derive summary statistics like a directed and undirected mean of the vector length, a standard deviation of the undirected vector length and a mean direction as an angle in degrees.

The many mean mobility vectors $\overrightarrow{x_{S_1}}, ..., \overrightarrow{x_{S_n}}$ are interesting observations for the individual samples $S_1, ..., S_i$. But they are not yet regional, diachronic mobility proxies. To achieve this, and to ultimately arrive at the grey mobility proxy curves in Figure 6, the sample-wise measurements have to be spatially and temporally binned and summarised. For spatial binning we defined non-overlapping subregions, for the time-segments we defined a heavily overlapping sequence of moving windows. For each space-time-bin we calculated the directed and undirected mean vector length, the standard deviation and standard error of the undirected mean vector length and again a mean direction angle.

### 3 Supplementary Text 3: Origin search parameter exploration

The mobility estimation presented in this paper depends on a large number of parameters. For many of them there is no naturally optimal choice, so we had to make multiple intuitive or empirically informed decisions. In the following sections we show and comment the effect of some alternative parameter choices on the final result (as displayed in Figure 6).

#### 3.1 MDS result dimensions

The multidimensional scaling analysis presented in the main text to obtain simplified ancestry components is deliberately limited to two result dimensions. A 2D "genetic map" is relatively easy to visualize and understand – and we could intuitively conclude that its structure is meaningful on the spatial and temporal scale of our analysis. We could generally rule out batch effects (e.g. systematic differences introduced by the data production in different labs). On the other hand, adding more dimensions to the MDS could potentially improve the model, as increasing the resolution could yield more precise results for the origin search. On a conceptional level, the three-dimensional, spatiotemporal space where human history played out may be represented best by a corresponding, three-dimensional genetic space. And a very cursory look at the empirical data seems to support this:

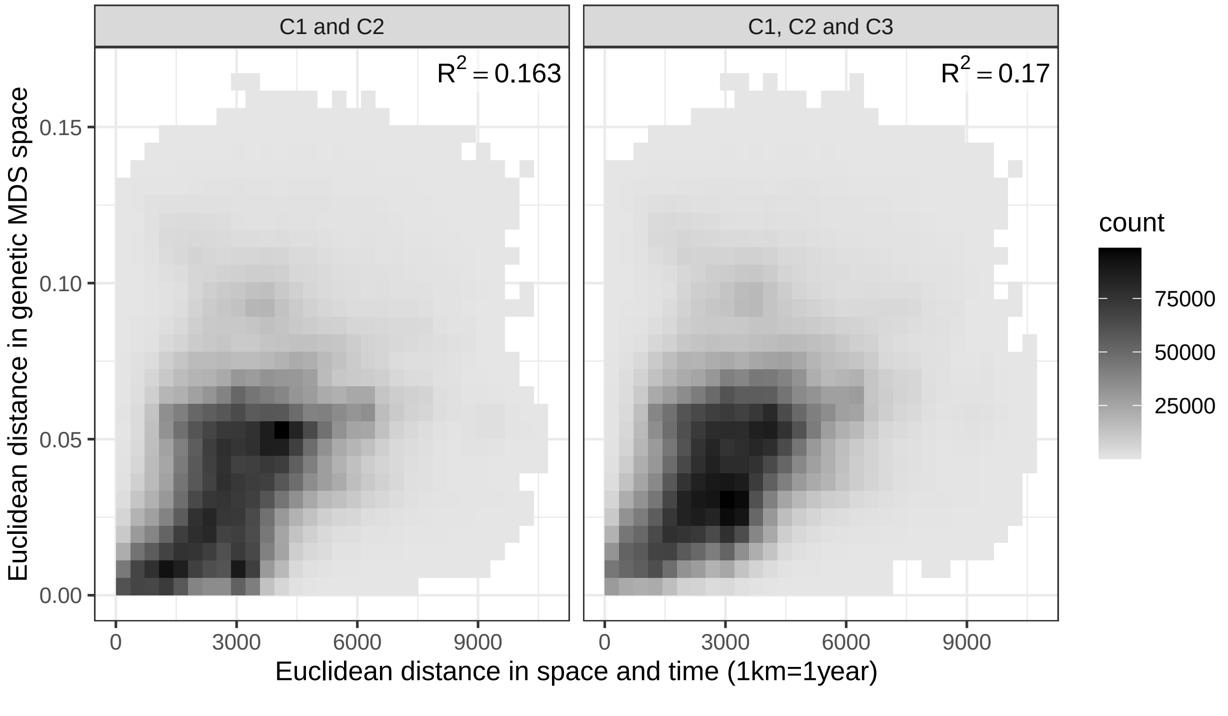


Supp. Figure XV: Correlation of pairwise genetic and spatiotemporal distance with genetic distance in two- or three-dimensional MDS space. The pairwise distances are counted in bins and plotted as a density raster.

Supp. Figure XV shows how an extremely simplified spatiotemporal distance (Euclidean spatiotemporal distance with 1 year = 1 kilometre) is correlated with Euclidean "genetic" distance in MDS space. Adding a third output dimension to the MDS seems to improve the correlation slightly, which is an argument to trust the third MDS dimension to represent some relevant and valuable information.

We went ahead and attempted to determine its nugget from the semivariogram, just as introduced above for Supp. Figure IV.

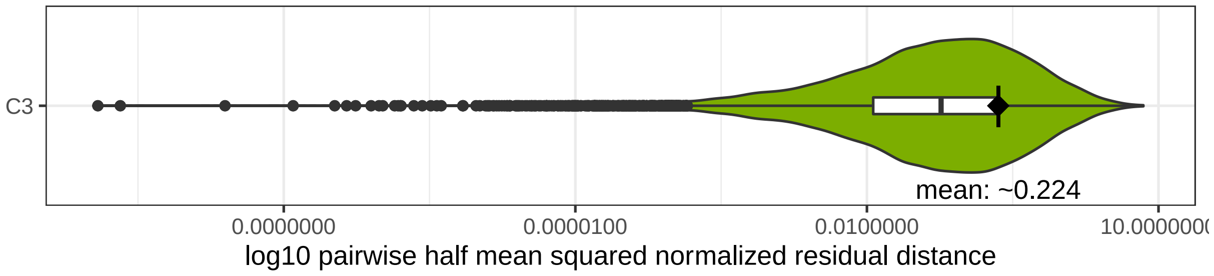


Supp. Figure XVI: Nugget estimation for the third MDS dimension. As Supp. Figure IV.

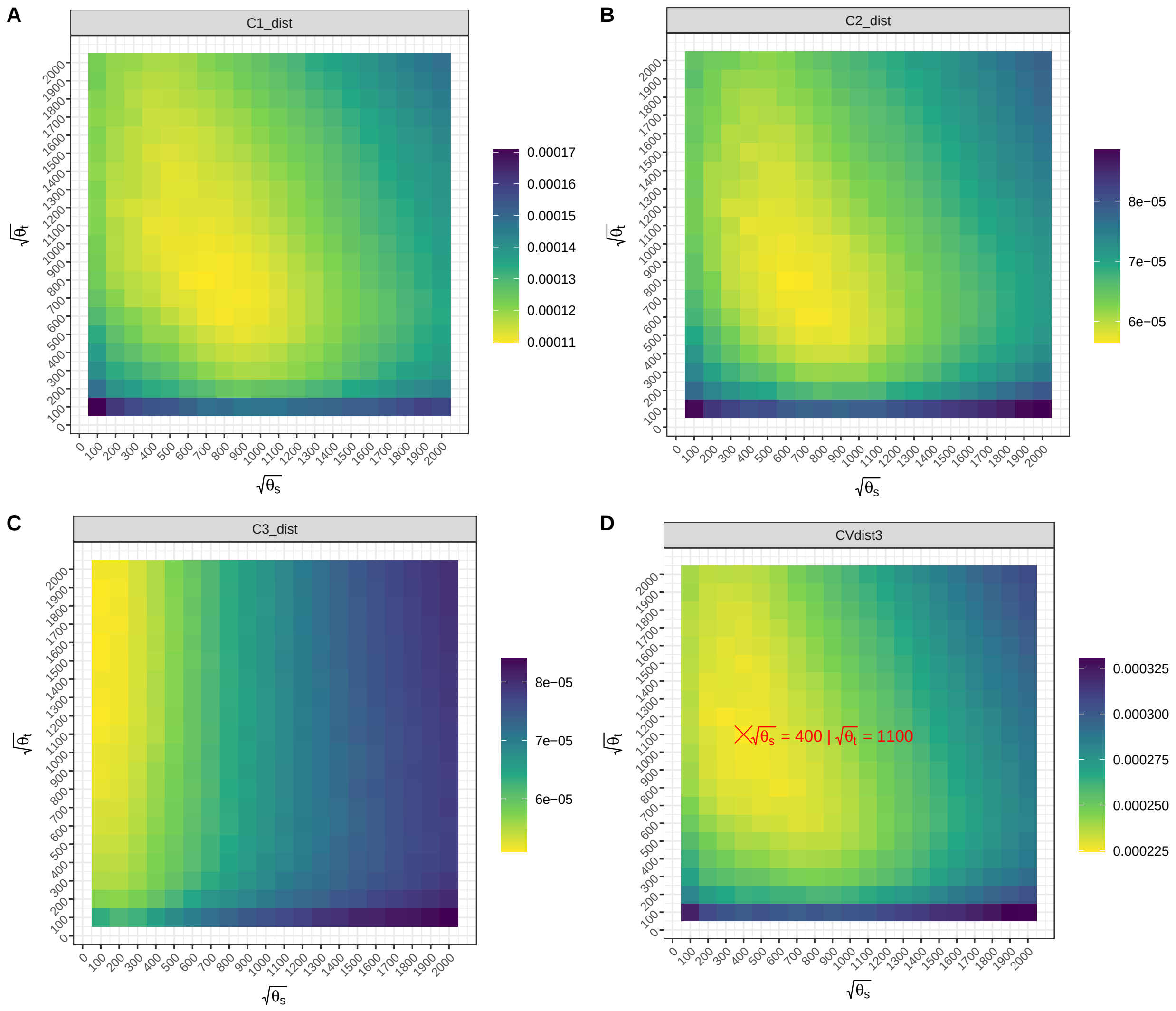


Supp. Figure XVII: Crossvalidation results as in Supp. Figure IX. The Nugget was fixed to 0.22 for this run.

The comparatively large nugget estimate highlighted in Supp. Figure XVI contradicts the previous observation and casts doubt on the third MDS dimension's consistency with a spatiotemporal signal. Kernel parameter estimation via crossvalidation confirms that C3 behaves differently to the generally similar C1 and C2 (Supp. Figure XVII).

Note that the nugget for this analysis was increased for all dependent variables, so not just C3, but also C1 and C2. Despite this change, the latter two yielded similar optimal kernel size results compared to what we saw in Supp. Figure IX above. C3, on the other hand, seems to be best postdicted by a GPR model with a large temporal, but extremely small spatial kernel, which is not well compatible with the other variables. See Supp. Table I for the parameters ultimately selected. The change in $\sqrt{\theta_t}$ also results in a different retrospection distance (Supp. Fig XIX).

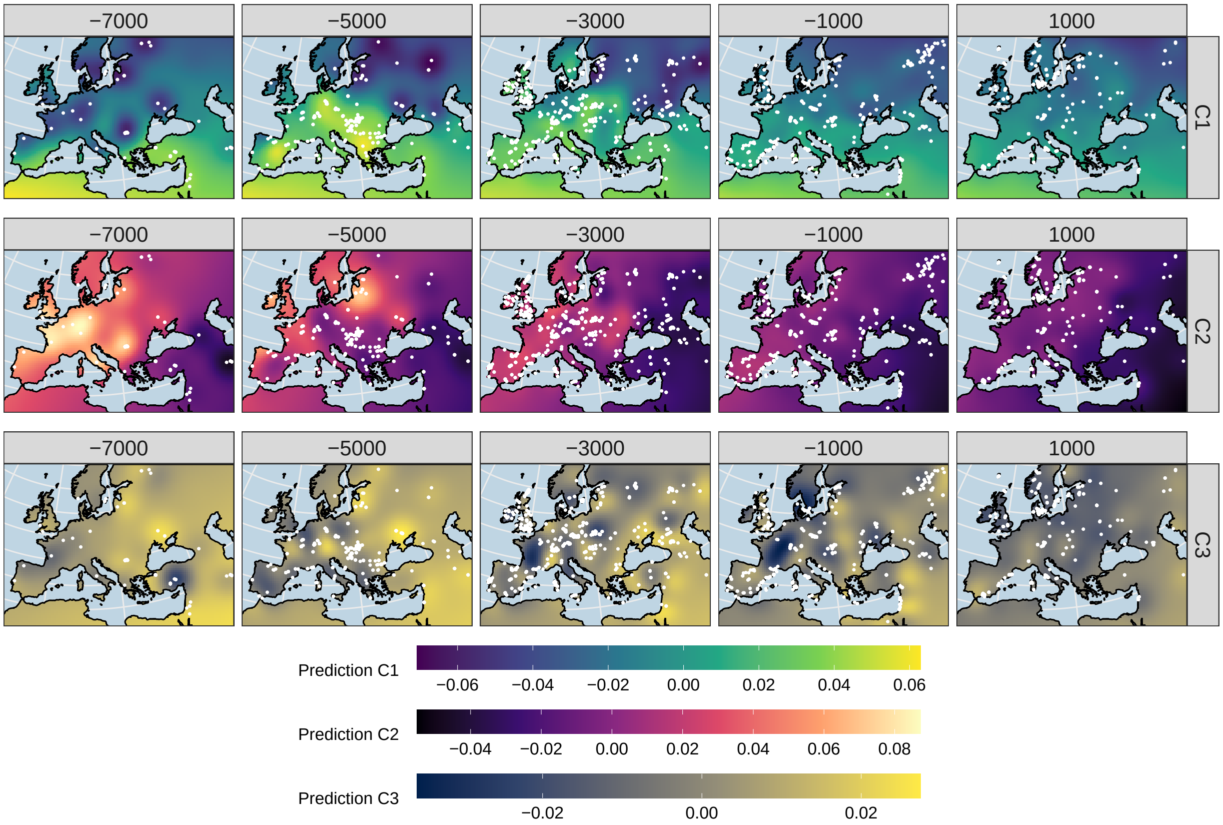


Supp. Figure XVIII: Gaussian process regression interpolation map matrix as in Figure 3, but here including the third MDS dimension.

Supp. Figure XVIII shows how the smaller spatial kernel size causes an even more pronounced "bull's eye effect" around individual observations, rendering the model theoretically more precise in regions and time periods with good data coverage, but also less accurate in regions and times where this is not the case. Finally we ran the origin search algorithm with this model to produce an equivalent of Figure 6, which we will compare below: Supp. Figure 1.

#### 3.2 Retrospection distance

For the retrospection/rearview distance in the origin search algorithm we decided to use an intermediate value based on the point of $Cov(x, x') = 0.5$ on the temporal dimension. We also tried two other settings: $Cov(x, x') = 0.75$ (Supp. Figure 2) and $Cov(x, x') = 0.25$ (Supp. Figure 3) years. See Supp. Table I and

Supp. Figure XIX for an overview of the parameters for these runs.

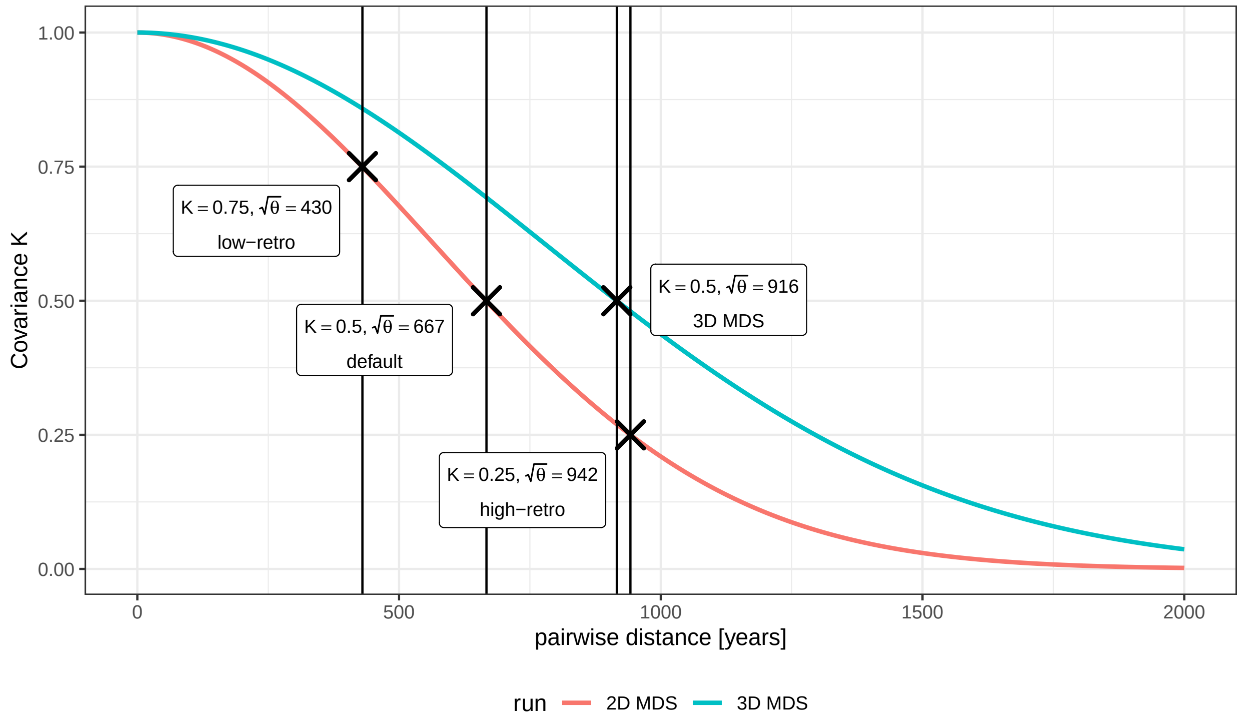


Supp. Figure XIX: Retrospection distance settings for the different runs. See Supp. Table I for a general overview of run parameters.

#### 3.3 Comparison

The curves for all three experimental settings (Supp. Figure 1, 2 and 3) are generally similar to the default that we selected for the main text (Figure 6). The main peaks and depressions generally overlap and seem to be detected robustly. There are a number of differences, though, especially visible when we calculate the divergence of individual-wise mobility vector means (Supp. Figure XX).

Longer retrospection distances generally cause longer mean mobility vectors. That is not surprising: It is plausible that the further one goes back in time, the further away ones ancestors might have lived originally. As the 3D MDS run relies on a field with a very large $\sqrt{\theta_t}$, which then also translates to a large retrospection distance, we assume the outcome of this run to be much more affected by this parameter than by any specific influence of the third dimension in genetic space.

An interesting deviation from the positive retrospection-mobility correlation emerges for Mesolithic individuals from Southeastern Europe (e.g. M95), who have a very local profile in the 3D MDS run. Ultimately we can not judge if this is a more or less correct representation of reality than the connection to Central Europe established in the other runs, but given the low amount of data for the latter region, we ascribed this observation in the main text to a methodological issue. Another interesting case are individuals from Ukraine from around 5000BC with a strong Eastern Hunter-Gatherer profile. They follow exactly the opposite trend than almost every other sample, as they appear significantly more non-local if one decreases the retrospection distance. This is probably a complex interaction of missing data and a thus badly informed ancestry field in

Scandinavia and the Forest Steppe region (see Figure 1).

The outlier individuals highlighted in the main text generally also appear as outliers in the alternative runs. There are some individuals, though, where this does not hold true. For example WEZ35 from the Late Bronze Age Tollense battlefield, which gets assigned much shorter mean mobility in the 3D MDS and the low retrospection distance run. The Early Neolithic individual I4971 from Hungary appears much more local with higher retrospection distances. LugarCanto45, I4064 and LON003 from Neolithic contexts in Portugal and Italy are attributed shorter mobility vectors in the 3D MDS run. I7807, finally, from Early Bronze Age Sicily, appears to be comparatively local in the low retrospection distance run – just as many other individuals from Italy around 2000BC. If Steppe ancestry spread into Italy in a wave-of-advance-like pattern, than lower mobility vectors for low retrospection distances are to be expected.

One last and special case we observed in the comparison of different runs, is a group of Middle and Late Neolithic (specifically 2880–2776calBC) individuals linked to the Globular Amphora culture in modern day Poland [4] (so in our analysis region Central Europe). They fell victim to violent conflict and were buried in a mass-grave at a time when Steppe-related ancestry was arriving. Genetically they can be clearly distinguished from the neighbouring Corded Ware groups, as they lack this new ancestry component. A conservative assumption would be, that these Globular Amphora culture individuals were a local group, not linked to any extreme long-distance mobility events in their recent past – unlike the Corded Ware newcomers from the east. Nevertheless these individuals, e.g. RISE1163 [4], show an extremely strong mobility signal with western direction in the 3D MDS run. The Globular Amphora culture is traditionally not thought to have emerged with an influx of western origin – on the contrary the scientific discourse revolves around an earlier potential connection to the east, which might have been relevant in its formation [5]. We believe the misleading signal in this specific run to be caused by a pressing-out effect, where the local, Central European ancestry profile in the interpolation field is significantly, yet erroneously modified by the Steppe ancestry of the newcomers. Three components are coming together here, as $\sqrt{\theta_t}$ for the 3D MDS run, the number of newly arriving individuals and also the distance in MDS space are all high in their respective frames of reference, thus amplifying the impact on the field. So for some of the last individuals with the "true" local ancestry profile – which happen to be the individuals from the Schroeder et al. paper – the origin search algorithm does not find the best correspondence in the local, skewed field, but 2000 kilometres further west.

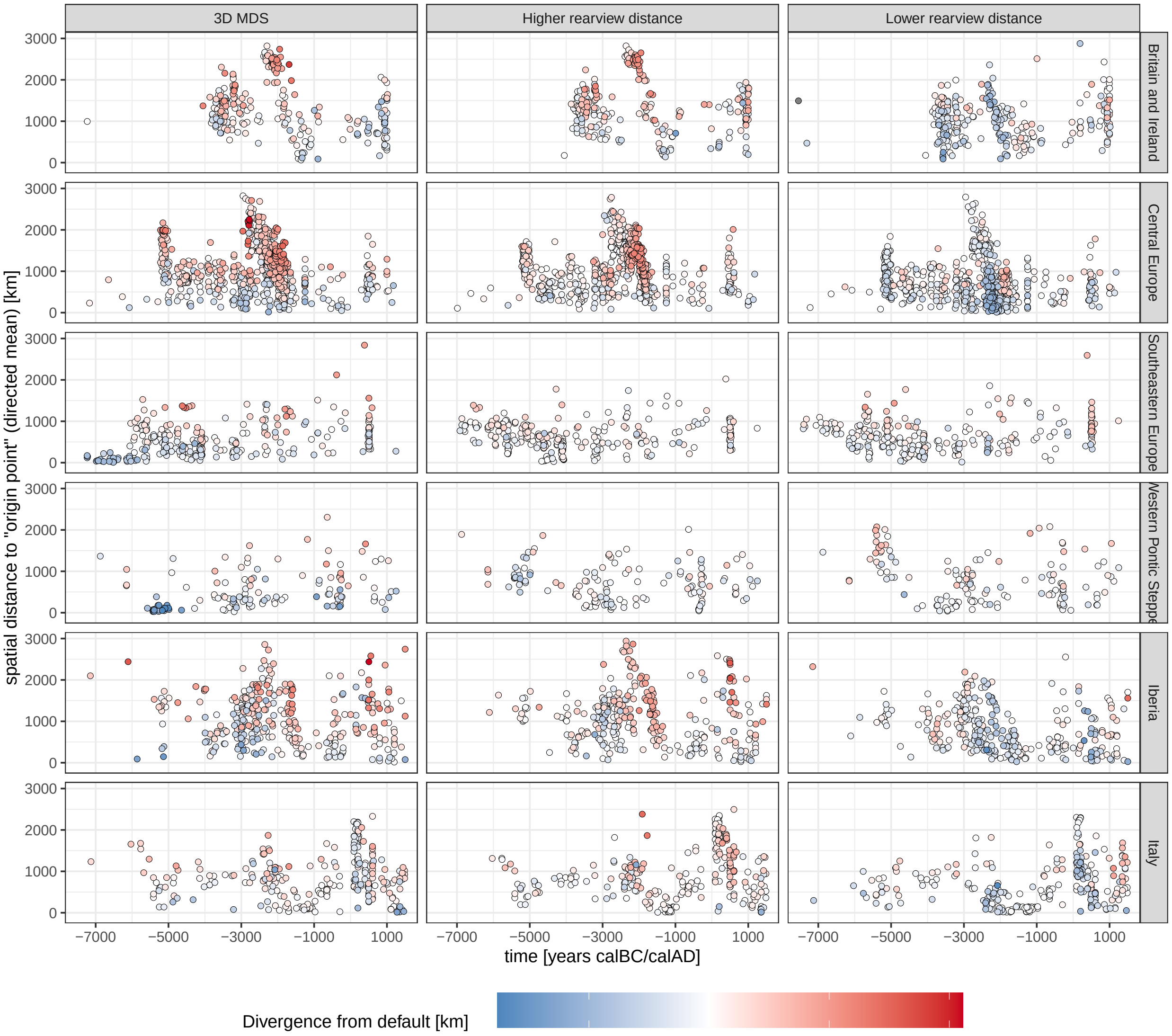


Supp. Figure XX: Comparison of different mobility estimation runs in a plot matrix. Each row of the plot matrix covers one analysis region, each column one of the three additional run configurations. The dots summarise the directed mean of all mobility vectors for one individual, just as the crosses in Figure 6. The diverging colour scheme shows if and how the different runs estimate mobility differently from the default run in the main text. Missing observations at the beginning of some of the sequences are due to long retrospection distances beyond the range of the interpolated field.

### 4 Supplementary Tables

Supplementary Table 1 lists all individuals/samples that went into the analysis and – only for the individuals from the regions that went through the origin search pipeline presented in the main text – summary statistics for said origin search. Here is a list with descriptions for the variables in this table:

1. **Individual_ID**: An identifier for the individual/sample (taken from the AADR's "Version ID")
2. **Group_Name**: A "population"/group the individual is attributed to in the AADR dataset (AADR: "Group ID")
3. **Country**: The modern day country the sample is coming from
4. **Region**: The spatial macroregion (as defined for this paper, see Figure 1) the sample is coming from
5. **Publication**: Publication from which the data for the respective sample was taken. The short publication keys are resolved on the AADR website: https://reich.hms.harvard.edu/allen-ancient-dna-resource-aadr-downloadable-genotypes-present-day-and-ancient-dna-data
6. **Search_x**: The spatial x-axis coordinate of the (archaeological) site a sample is coming from. Coordinates are given in metres according to EPSG:3035 (ETRS89 Lambert Azimuthal Equal-Area, "European grid") and rounded to kilometres (-3 decimal places) after conversion from the AADR's WGS 84 lat-lon coordinates
7. **Search_y**: The respective y-axis coordinate
8. **Search_z**: The (post-radiocarbon calibration) median age of the sample in years calBC/AD (negative values are BC). As the origin search was repeated in many iterations with different ages drawn from the post-calibration probability distribution, this median age only stands representative for the actual ages used for individual runs
9. **Search_C1**: The first output coordinate of the multidimensional scaling analysis for this sample. All MDS coordinates are rounded to four decimal places
10. **Search_C2**: The second MDS output coordinate
11. **Search_C3**: The third MDS output coordinate. This is given here, despite it was not considered for the origin search run summarised in the following columns of this table (see Supplementary Text 3)
12. **Origin_x**: The spatial x-axis coordinate of the centroid for all positions found with the default origin search algorithm: Each temporal resampling run yields one output position, and here we list the mean position for all of them
13. **Origin_y**: The respective y-axis coordinate
14. **Origin_z**: The mean age of the ancestry field time slices in which the origin search was performed. The different input ages (see Search_z) have the consequence, that the origin search is also performed in different time slices. We summarise their age as a simple mean here, rounded to decades
15. **Origin_C1**: The mean value of the first output coordinate in MDS space as derived from the interpolated ancestry field for all minimum distance points across the origin search iterations. This value will be close, but generally not identical to Search_C1 (see Supplementary Text 2)

16. **Origin_C2**: The equivalent for the second MDS coordinate

17. **Undirected_mean_spatial_distance**: The mean of all distances from the sample to its potential origin points across all temporal resampling iterations. This mean is *undirected* in the sense that it works on the absolute distances, so only the lengths of the origin vectors connecting search and origin point. Rounded to kilometres

18. **Directed_mean_spatial_distance**: This mean distance considers the direction of the origin vectors, so vectors with opposing directions cancel each other out. It is therefore smaller than the undirected one. Rounded to kilometres

19. **Mean_angle**: Mean direction of the origin vectors as an angle in degree ($0 - 360°$). Rounded to full degrees

20. **Mean_cardinal_direction**: Translation of the mean angle to a simple cardinal direction (N, E, S, W)
